# Decoding cellular transcriptional regulatory networks governing wheat inflorescence development

**DOI:** 10.1101/2025.01.18.633750

**Authors:** Xuemei Liu, Xuelei Lin, Jingmin Kang, Katie A. Long, Jingjing Yue, Chuan Chen, Dongzhi Wang, Ashleigh Lister, Iain C. Macaulay, Xin Liu, Cristobal Uauy, Jun Xiao

## Abstract

Wheat inflorescence architecture, particularly spikelet and floret development, is critical for grain yield. To decode the cellular transcriptional regulatory network (cTRN) underlying wheat inflorescence development, we integrated multiple single-cell omics technologies to construct a spatiotemporal atlas of transcriptional and chromatin accessibility dynamics. This comprehensive analysis identified 20 cell types, 7,211 cell type-specific genes and 152,333 cell type-specific accessible chromatin regions (csACRs) in the wheat inflorescence. Trajectory analysis identified two sub-clusters of proliferating cells as the origins of spikelet and floret formation, deviating from the traditional developmental model. Key transcription factors and hormone-related genes in the cTRN, along with the csACRs, providing new targets for modulating wheat inflorescence architecture. Our findings provide a high-resolution resource for crop inflorescence research and establish a paradigm for applying spatiotemporal single-cell omics analysis to plant biology.

## Introduction

Inflorescence architecture is crucial for crop yield and fitness. The spatial arrangement and number of flowers, which are tightly regulated by genetic factors, directly influence the number and size of seeds per inflorescence. In cereals like wheat (*Triticum aestivum*), flowers (termed florets) develop within spikelets attached to the inflorescence axis in a distichous phyllotaxis (Whipple 2017; Koppolu et al. 2022). This intricate architecture is established when the vegetative meristem transitions into an inflorescence meristem (IM), marking the start of the double ridge (DR) stage. During this stage, the IM produces two ridges: the upper ridge developed into spikelet meristems (SM), while the lower ridge produces leaf primordia which are actively repressed (Koppolu and Schnurbusch 2019; Luo et al. 2023). The IM continues generating double ridges until the formation of the terminal spikelet (TS) which determines the final number of spikelets (Sakuma and Schnurbusch 2020; Wang et al. 2021). As development proceeds, the SM sequentially produce leaf-like structures (glumes and lemmas) and floret primordia (FP), which develop into floral organs and grains. Unlike the determinate IM, the SM remains indeterminate, typically initiating over 12 florets (Sakuma and Schnurbusch 2020; Luo et al. 2023), though only 3-5 florets remain fertile and produce grains (Shitsukawa et al. 2009).

Numerous genes regulate inflorescence architecture by controlling spikelet and floret patterning, as well as their number, during inflorescence development (Reviewed by (Koppolu and Schnurbusch 2019; Yoshikawa and Boden 2024). Among these, MADS-box transcription factors (TFs) are central to this regulation, particularly in spikelet and floret development. For example, ectopic expression of *SHORT VEGETATIVE PHASE-2* (*SVP2*) results in elongated organs within spikelets and increased rudimentary basal spikelets (Adamski et al. 2021; Liu et al. 2021), while *SEPALLATA* (*SEP*) E-class genes are essential for normal floral development (Li et al. 2021a). TFs from the APETALA2/ETHYLENE RESPONSE FACTOR (AP2/ERF) family, such as *FRIZZY PANICLE* (*WFZP*) and *DUO*, cause supernumerary spikelet (SS) phenotypes when mutated (Dobrovolskaya et al. 2015; Du et al. 2021; Li et al. 2021b; Wang et al. 2022).

Plant hormones also play critical roles in inflorescence development (Taylor-Teeples et al. 2016; Cucinotta et al. 2021). In *Arabidopsis*, rice (*Oryza sativa*), and maize (*Zea mays*), local auxin maxima, mediated by the auxin transporter PIN FORMED 1 (PIN1), initiate various primordium formation within the inflorescence (Benková et al. 2003; Gallavotti et al. 2008; Yang et al. 2017). Loss of PIN1 in *Arabidopsis* results in pin-shaped inflorescence with no floret initiation (Reinhardt et al. 2000). Cytokinin levels also affect inflorescence architecture, with mutations in the cytokinin metabolism gene *CYTOKININ OXIDASE/DEHYDROGENASE 2* (*OsCKX2*) increasing spikelet numbers and yield in rice (Ashikari et al. 2005). Additionally, changes in brassinosteroids-related genes (BRs) impact spikelet formation, with altered BR signaling in the secondary branch meristem enhancing grain number in rice (Zhang et al. 2024b).

While the effects of these and other individual genes on wheat inflorescence development are increasingly understood (Gao et al. 2019; Luo et al. 2023), the molecular networks governing spikelet and floret formation remain incomplete. This knowledge gap is partly due to most networks relying on whole inflorescence samples, averaging across multiple tissues and cell types (Li et al. 2018; Wang et al. 2022; Lin et al. 2024). Such bulk approaches fail to capture critical transcriptional differences arising from asynchronous development patterns among spikelets (Backhaus et al. 2022; Long et al. 2024). Furthermore, isolating single spikelets or florets, is technically challenging particularly during early stages.

Recent advancements in single-cell omics techniques, including single-nucleus RNA sequencing (snRNA-seq; (Zhang et al. 2021a; Zong et al. 2022), snATAC-seq (Marand et al. 2021; Zhang et al. 2024a), and Stereo-seq (Xia et al. 2022; Zhang et al. 2024a), provide powerful tools to dissect plant developmental processes. Single-cell expression analyses uncover gene functional redundancies that are often obscured in bulk data. For instance, in maize branching mutant *ramosa3* (*ra3*), both *TREHALOSE PHOSPHATE PHOSPHATASE 4* (*ZmTPP4*) and *ZmTPP12* are upregulated, but only *ZmTPP4* shows redundancy with *RA3* at the cellular level, despite bulk RNA-seq co-expression (Xu et al. 2021). Spatial transcriptomics, such as Stereo-Seq, complements snRNA-seq by distinguishing cell types within complex tissues. For example, Stereo-seq successfully classified cell types in the maize ear meristem into indeterminate and determinate types (Wang et al. 2024). Moreover, single-nucleus assay for transposase-accessible chromatin with sequencing (snATAC-seq) provides high resolution of gene regulatory networks, identifying tissue- or cell type-specific genes and cis-regulatory elements (CREs) (Zhang et al. 2024a). The insights gained from these methods are crucial for optimizing inflorescence architecture to increase yield potential, as emerging evidence suggests that precise control of spatiotemporal gene expression could help reconcile trade-offs between various traits like grains per panicle versus tiller number (Song et al. 2022), or grain size versus grain number in rice (Zhang et al. 2024b).

In this study, we integrated snRNA-seq, snATAC-seq, and Stereo-seq to map the spatiotemporal transcriptional and chromatin dynamics of wheat inflorescence development. We identified 20 cell types, 7,211 cell type-specific genes, and 152,333 csACRs, providing a high-resolution resource for gene functional studies. Trajectory analysis uncovered proliferating cell sub-clusters involved in spikelet and floret formation. The identified cellular transcriptional regulatory network (cTRNs) and csACRs offer novel targets for modulating wheat inflorescence architecture and improving yield potential.

## Results

### Cellular gene expression and accessible chromatin atlas of developing wheat inflorescence

To unravel the cellular gene expression regulation during wheat inflorescence development, we employed single-cell omics techniques, generating 12 snRNA-seq and 20 snATAC-seq libraries spanning six developmental stages, from late double ridge (stage W2.5) to floral organ formation (W5) (WADDINGTON et al. 1983) (Fig. 1A). The average unique molecular identifier (UMI) count across snRNA-seq libraries ranged from 3,200 to 10,000, covering 2,500 to 5,000 genes per cell (Fig. S1A). For snATAC-seq, the UMI count also ranged from 3,200 to 32,000, with the fraction of reads in peaks (FRiP) values exceeding 0.4 across libraries (Fig. S1B, D, E). After rigorous quality control, we retained 73,729 cells from snRNA-seq and 75,745 cells from snATAC-seq (Fig. S1A-E, Table S1, S2). The data demonstrated high reproducibility across biological replicates (Fig.1B, Fig. S1C) and reflected patterns observed in bulk ATAC-seq previously (Lin et al. 2024) (Fig. S1F).

**Fig. 1.**
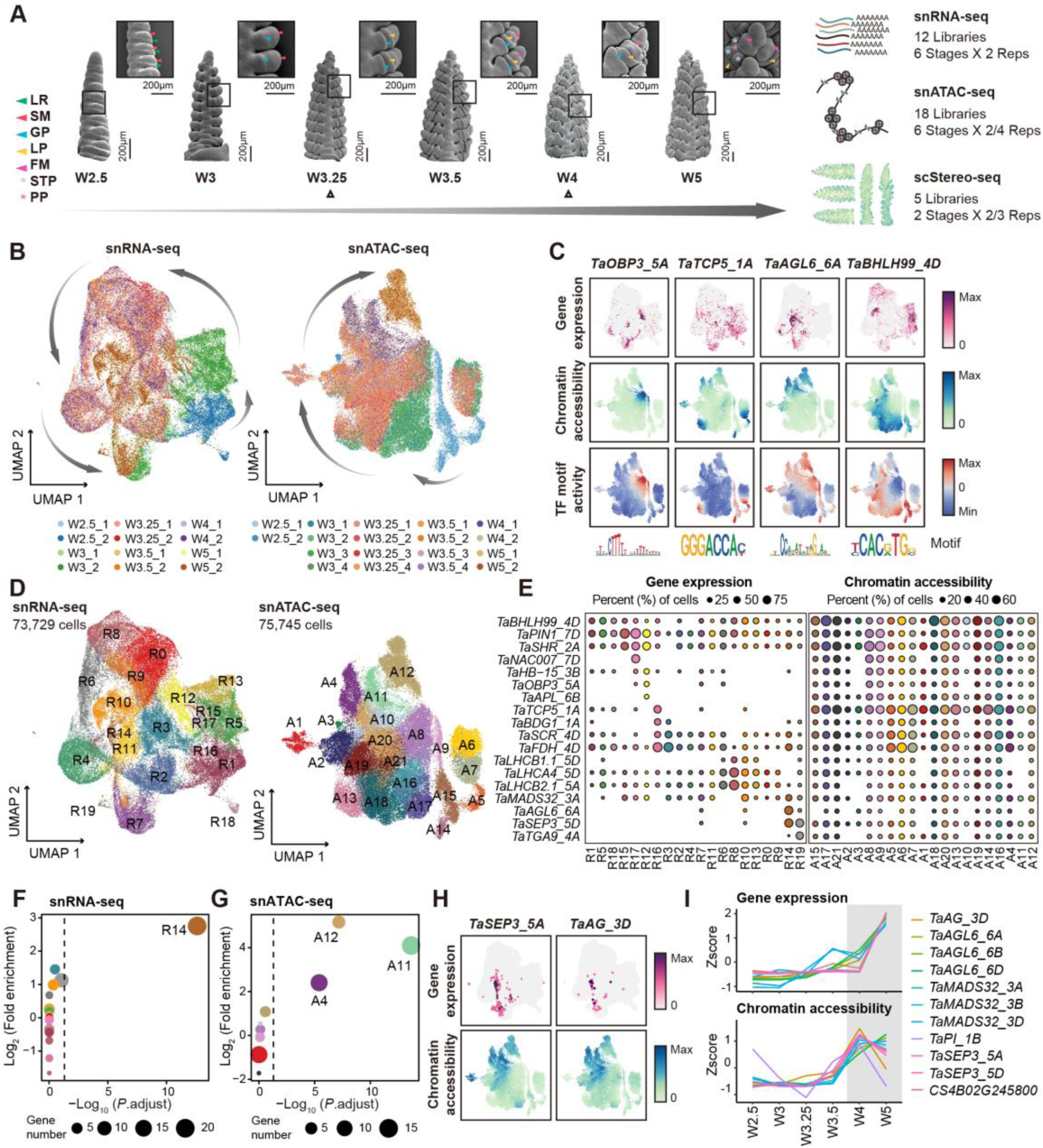
A single-cell transcriptomic and chromatin atlas of wheat inflorescence. **A** Experimental design. Wheat inflorescence morphology was observed via scanning electron microscopy at six developmental stages. LR, leaf ridge; SM, spikelet meristem; GP, glume primordium; LP, lemma primordium; FM, floret meristem; STP, stamen primordium; PP, pistil primordium. The triangle marks the two stages during which the scStereo-seq experiment was conducted. **B** UMAP visualization of cells, colored by different snRNA-seq (left) and snATAC-seq (right) libraries. **C** Profiles of gene expression (snRNA-seq), chromatin accessibility (snATAC-seq) and TF motif activity (snATAC-seq) for *TaOBP3_5A*, *TaTCP5_1A*, *TaAGL6_6A* and *TaBHLH99_4D*. **D** UMAP visualization of cells, colored by snRNA-seq (left) and snATAC-seq (right) cell types. **E** Dotplot showing the percentage of cells expressing selected genes across snRNA-seq cell types (left) and the percentage of cells with open chromatin of selected genes across snATAC-seq cell types (right). Dots with a proportion below 5% are not displayed. **F** Enrichment of MIKC_MADS family genes in cell type-specific genes across different cell types in snRNA-seq. **G** Enrichment of MIKC_MADS family genes in cell type-specific genes across different cell types in snATAC-seq. **H** Gene expression and chromatin accessibility profiles of *TaSEP3_5A* and *TaAG_3D*. **I** Z-scaled gene expression and chromatin accessibility levels of MIKC_MADS genes across different developmental stages.

Cells from different developmental stages exhibited substantial heterogeneity, following a continuous developmental trajectory. We integrated the datasets into a unified single-cell map depicting inflorescence development (Fig. 1B). Notably, TFs from families such as TCP, bHLH, and MADS exhibited cell type-specific expression patterns, and their chromatin accessibility and TF binding motif activity were elevated in specific cell types, suggesting coordinated regulation of gene expression (Fig. 1C). Additionally, subgenomic expression asymmetry was higher at the single-cell level (35.5% balanced triads) compared to bulk RNA-seq (73.3%) from spikes (Ramírez-González et al. 2018), consistent with previous findings in root tissue (Zhang et al. 2023) (Fig. S1G).

Using uniform manifold approximation and projection (UMAP) for dimensionality reduction followed by clustering analysis, we identified 20 distinct cell types in snRNA-seq (R0 to R19) and 21 in snATAC-seq (A1 to A21) (Fig. 1D). Among these, 7,211 and 18,698 cell-type-specific genes were identified in the snRNA-seq and snATAC-seq data, respectively (Table S3, S4). Only 920 genes were dual-cell-type specific for both expression and chromatin accessibility, with transcription showing greater specificity than chromatin accessibility (Fig. 1E). MADS TFs play a key role in inflorescence development (Hama et al. 2004; Shitsukawa et al. 2009; Kong et al. 2022). Our analysis revealed MADS TFs were enriched in cell type-specific genes within R14 (snRNA-seq) and A4, A11, A12 (snATAC-seq) (Fig. 1F, G), exemplified by genes like *AGAMOUS-LIKE 6* (*TaAGL6_6A*), *TaSEP3_5A* and *AGAMOUS* (*TaAG_3D*) (Fig. 1C, H). suggesting a potential correspondence between these cell types. However, they showed different developmental timing: R14 predominated at W5, while A4 and A11 were observed at W4, with A12 at W5 (Fig. 1B, D). This distribution indicates that chromatin opening of MADS genes precedes peak gene expression (Fig. 1I), highlighting a temporal ‘priming’ pattern.

These results highlight the power of snRNA-seq and snATAC-seq in capturing cellular heterogeneity and temporal dynamics, offering a rich resource for exploring wheat inflorescence development at cellular resolution.

### scStereo-seq and MERFISH provide spatial context for cell types annotation

To better understand the complex cellular architecture of wheat inflorescence, we used spatial transcriptomic techniques to annotate cell types identified by snRNA-seq and their gene expression (Fig. S2A). We conducted scStereo-seq on wheat inflorescences from the W3.25 and W4 stages (Fig. S2B), yielding 20,183 high-quality cells (Fig. S2C, D, Table S5, See methods for details). The data showed no positional bias in gene expression or cell count (Fig. S2E, F), and demonstrated high repeatability across sections from the same stage (Fig. S2G). To map snRNA-seq data to spatial gene expression, we used Tangram to align gene expression based on shared genes (Fig. S2A). Combining cell type localization (Table S5), gene expression patterns (Fig, 2A-C, Fig. S3), TF family analysis, and gene ontology (GO) enrichment (Fig. 2E, F, Table S6, S7) with the multiplexed error-robust fluorescence *in situ* hybridization (MERFISH) data (Long et al. 2024) (Fig. 2H), we categorized wheat inflorescence cell types into three distinct groups (Fig. 2D).

**Fig. 2.**
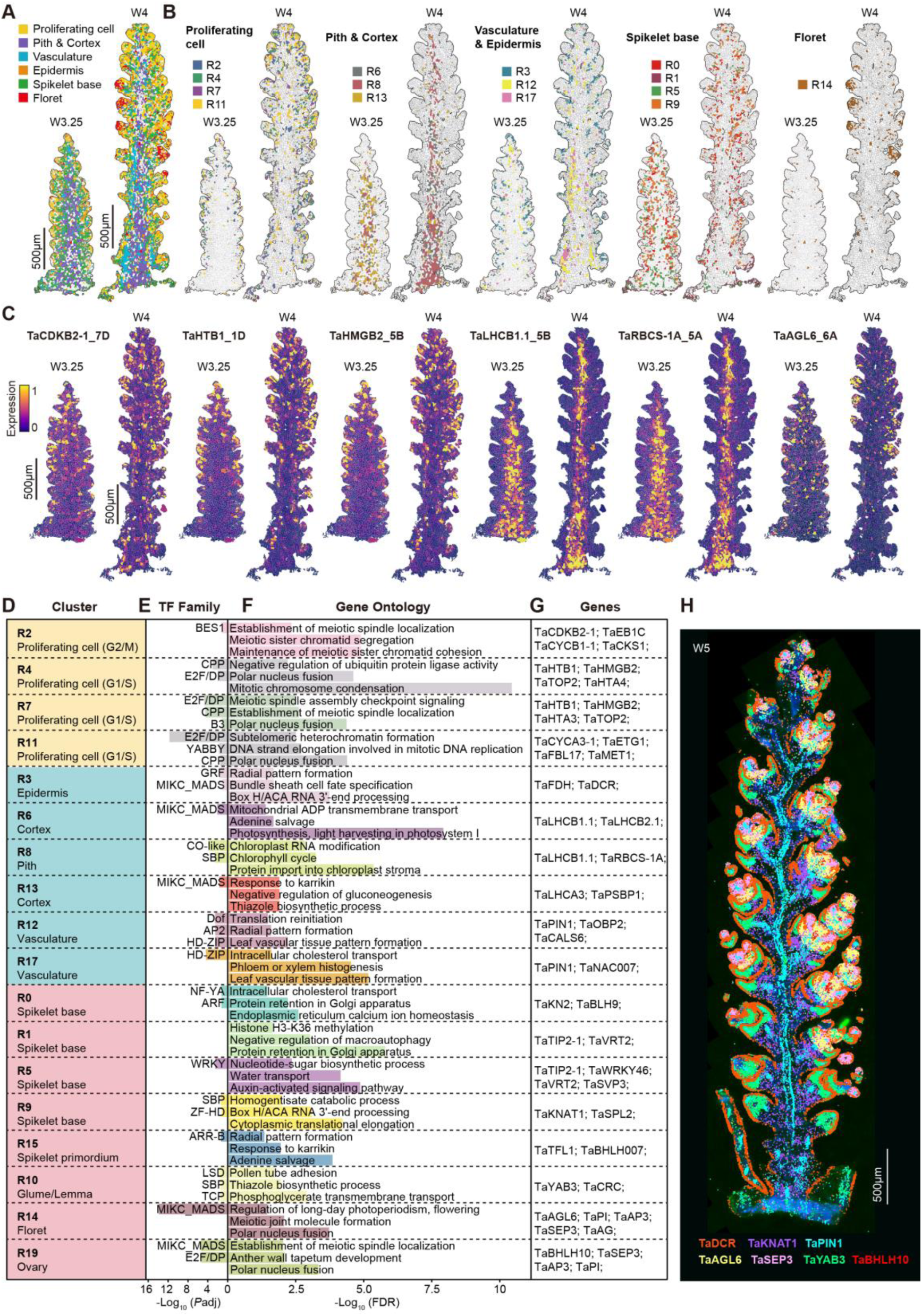
Cell type annotation for developing wheat inflorescence. **A, B** Spatial distribution of cell types defined by snRNA-seq mapped on scStereo-seq data. **C** Predicted spatial gene expression of cell type-specific genes by mapping the snRNA-seq data to scStereo-seq data. **D** Cell type annotation results. **E** TF family enrichment analysis for cell type-specific genes. The top three enriched TF families are shown (two-sided Fisher’s exact test, Benjamini and Hochberg (BH) correction for multiple testing, padj < 0.05). **F** GO enrichment analysis for cell type-specific genes (two-sided Fisher’s exact test, False Discovery Rate (FDR) correction for multiple testing, FDR < 0.05). **G** Names of cell type-specific genes. **H** MERFISH of multiple cell type-specific genes during wheat inflorescence development. Different colored dots represent different genes.

The first category comprises proliferating cells, including R2, R4, R7, and R11, located in spikelets with active cell division (Fig. 2A, B, Fig. S3A, B). These cells express genes involved in mitosis and chromosome condensation, such as *CYCLIN DEPENDENT KINASE B2;1* (*TaCDKB2-1*) (Vandepoele et al. 2002), *HISTONE 2B 1* (*TaHTB1*) (Jiang et al. 2020); and *HIGH MOBILITY GROUP B2* (*TaHMGB2*) (Pedersen and Grasser 2010) (Fig. 2C-G, Fig. S3C, Fig. S4). These cell show marker genes specific to different cell cycle phases (Francis 2007), with *TaCDKB2-1* and *CYCLIN B1;1* (*TaCYCB1-1*) marking the G2/M phase in R2, while *TaCYCA3-1* and *E2F* family genes marking the G1/S phase in R4, R7, and R11 (Fig. 2D, G, Fig. S4).

The second category consists of structure cells providing support to the inflorescence, including epidermis (R3), vasculature (R12, R17), pith (R8), and cortex (R6, R13) (Fig. 2A, B, Fig. S3A, B). Marker genes like *FORMATE DEHYDROGENASE* (*FDH*) and *DEFECTIVE IN CUTICULAR RIDGES* (*DCR*), specific to *Arabidopsis* epidermis (Zhang et al. 2021a), are highly expressed in the epidermis (R3) (Fig. 2D, G, H, Fig. S4), while vasculature cells (R12, R17) express the auxin efflux carrier *TaPIN1* (Scarpella et al. 2006), essential for vascular tissue formation and phloem/xylem histogenesis (Fig. 2D-H, Fig. S4). Pith (R8) and Cortex (R6, R13) cells, located at the central axis, express genes like *LIGHT HARVESTING CHLOROPHYLL A/B-BINDING 1.1* (*TaLHCB1.1*) (Sawchuk et al. 2008) and *RIBULOSE BISPHOSPHATE CARBOXYLASE SMALL CHAIN 1A* (*TaRBCS-1A*) (Sawchuk et al. 2008), involved in photosynthesis and energy metabolism (Fig. 2A-G, Fig. S3C, Fig. S4).

The third category includes developmental cells responsible for spikelet and floret formation, such as spikelet primordium (R15), spikelet base (R0, R1, R5, R9), glume/lemma (R10), floret (R14), and ovary (R19) (Fig. 2A, B, Fig. S3A, B). Expression of *TERMINAL FLOWER 1* (*TaTFL1*) (Sun et al. 2023), and *BASIC HELIX-LOOP-HELIX 007* (*TaBHLH007*) (Zhang et al. 2021b) defines R15 as spikelet primordium (Fig. 2D, G, Fig. S4), while genes like *KNOTTED-LIKE FROM ARABIDOPSIS THALIANA 1* (*TaKNAT1*) (Douglas et al. 2002) and *TaKN2* (Postma-Haarsma et al. 2002) in R0, R1, R5, and R9 regulate inflorescence architecture (Fig. 2D, G, H, Fig. S4). R10 is marked by *YABBY3* (*TaYAB3*) expression in the glume and lemma, and R14 is involved in floret formation with expression of genes like *TaAGL6* and *TaSEP3* (Fig. 2A-D, G, H, Fig. S4). R19 cells are specific to ovary, marked by *TaBHLH10* expression (Fig. 2D, G, H, Fig. S4).

Spatial transcriptomics from scStereo-seq and MERFISH complement snRNA-seq, enabling detailed annotation of cell types during inflorescence development.

### Distinct trajectories of spikelet and floret formation from proliferating cell sub-clusters

Spikelet and floret initiation are crucial for grain number determination and yield (Sakuma and Schnurbusch 2020). To explore their developmental trajectories, we utilized Monocle3 (Cao et al. 2019) for trajectory inference based on annotated cell types. The cells were grouped into five partitions (P1-P5), with R15 (spikelet primordium) in P4 and R14 (floret) in P1 (Fig. 3A, B, Table S8). Notably, R7 (proliferating cells) was split into two sub-clusters, R7.P4 and R7.P1, belonging to P4 and P1, respectively (Fig. 3A, B). Seurat (Hao et al. 2021) analysis also distinguished these sub-clusters (Fig. 3C).

**Fig. 3.**
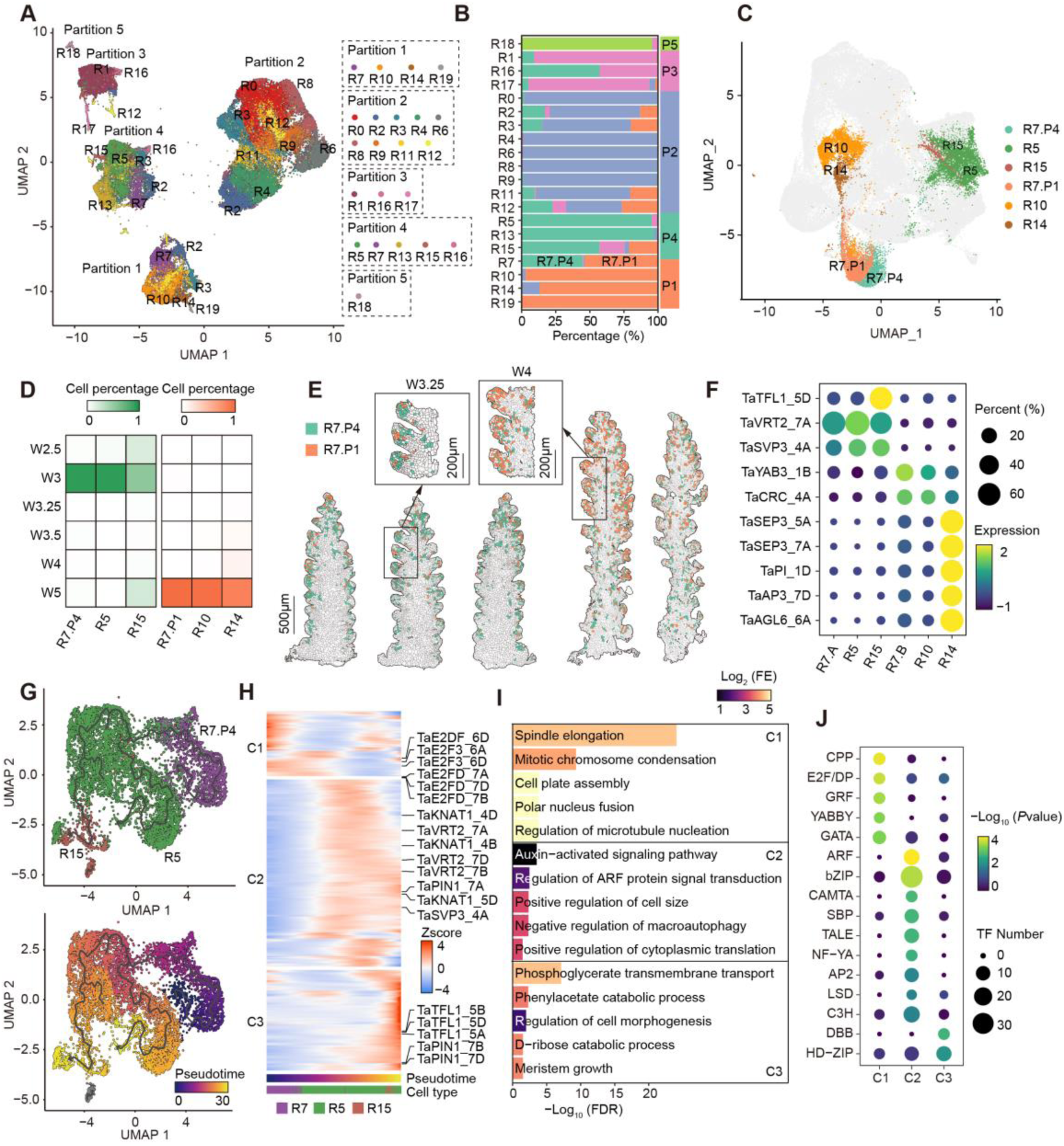
Trajectory inference for wheat spikelet and floret formation. **A** UMAP visualization of cells processed using Monocle3, colored according to snRNA-seq cell types. **B** Proportion of cells assigned into different partitions across various snRNA-seq cell types. **C** Seurat UMAP visualization of cells belonging to R5, R7.P1, R7.P4, R10, R14, and R15. **D** Proportion of cells at various developmental stages. **E** Spatial distribution of R7.P4 and R7.P1 cells in snRNA-seq mapped on scStereo-seq data. **F** Dotplot showing the gene expression level and the percentage of cells expressing DEGs of R7.P4 and R7.P1. **G** Trajectory inference and pseudotime analysis of cells belonging to R7, R5, and R15 in partition4. **H** Heatmap showing the clustering results of genes whose expression change across pseudotime of spikelet formation. **I** GO enrichment analysis of genes in different clusters during spikelet formation (two-sided Fisher’s exact test, FDR for multiple testing, FDR < 0.05). FE: fold enrichment. **J** TF family enrichment of genes in different clusters during spikelet formation (two-sided Fisher’s exact test, *P*value < 0.01).

Indeed, the two sub-clusters of R7 showed distinct spatiotemporal patterns; 95% of R7.P4 cells were present at W3 stage, corresponding to early spikelet development, while 99% of R7.P1 cells were present at W5 stage, aligning with floret development (Fig. 3D). Spatial mapping to scStereo-seq data further confirmed that R7.P4 cells were localized in the spikelet at W3.25, while R7.P1 cells were predominant within the developing meristem in florets at W4 (Fig. 3E). Differentially expressed genes (Table S9) revealed that *TaTFL1* and *VRT2*, associated with spikelet development (Adamski et al. 2021; Li et al. 2021a; Liu et al. 2021; Sun et al. 2023), were highly expressed in R7.P4, R5, and R15. Meanwhile, *TaSEP3*, involved in floral organ development (Paolacci et al. 2007), was upregulated in R7.P1, R10 and R14 (Fig. 3F). These findings suggest that R7.P4 and R7.P1 cells distinctly contribute to spikelet and floret formation.

To further investigate the transcriptional dynamics during spikelet and floret formation, we performed trajectory inference and pseudotime analysis on cells from R7.P4 (proliferating cells), R5 (spikelet base), and R15 (spikelet primordium) in partition P4, as well as R7.P1 (proliferating cells), R10 (glume/lemma), and R14 (floret) in partition P1 (Fig. 3G, S5A, Table S10, S11). The predicted trajectory revealed the formation process of R5 (spikelet base) and R15 (spikelet primordium) cells in partition P4 (Fig. 3G), and R10 (glume/lemma) and R14 (floret) cells in partition P1 (Fig. S5A).

We identified three clusters of genes exhibiting pseudotime-dependent expression changes (Moran’s I, q-value < 1e-10) in partition P4 (Fig. 3H, Table S12). Cluster C1 genes, involved in cell division, were enriched with E2F/DP family TFs like *TaE2FD* (Fig. 3H-J). Cluster C2 genes, associated with auxin signaling and cell size regulation, featured ARF, bZIP, SBP, and TALE family TFs, including the Class-I KNOTTED1-like homeobox (KNOX) gene *TaKNAT1*, which regulates inflorescence architecture (Fig. 3H-J) (Douglas et al. 2002). Cluster C3 genes, predominantly expressed in the spikelet primordium (R15), were linked to meristem growth and cell morphogenesis, enriched with HD-ZIP and bZIP TFs (Fig. 3H-J). For partition P1, pseudotime-dependent genes were grouped into three clusters (Fig. S5B, Table S13). Cluster G1 genes, associated with proliferation, included cell cycle-related E2F/DP TFs (Fig. S5B-D). Cluster G2 genes, expressed in the glume/lemma (R10), were involved in leaf morphogenesis and abaxial cell fate specification, exemplified by *TaYAB3* (Fig. S5B-D). Cluster G3 genes included MADS family members such as *TaPI*, *TaSEP3*, and *TaAG*, essential for specifying floret identity (R14) (Fig. S5B-D).

This comprehensive trajectory analysis highlights the dynamic processes by which spikelet and floret differentiate from two distinct sub-clusters of proliferating cells, enhancing our understanding of wheat inflorescence development at the cellular level.

### Cell type-specific ACRs and associated TFs guide spatiotemporal gene expression

Spatiotemporal gene expression is vital for wheat inflorescence development. To investigate its regulation, we integrated snRNA-seq and snATAC-seq data. Initially, we aligned cell types across both datasets, finding the highest overlap between A11-specific genes in snATAC-seq and R14 in snRNA-seq (19%), suggesting a connection between these cell types (Fig. 4A, Fig. S6A). Using a 15% overlap threshold and requiring at least 10 overlapping genes, we aligned other cell types between the datasets (Fig. S6A). Major cell types in snATAC-seq include epidermis (A4, A5, A6, A7), vasculature (A8, A9), pith (A10, A19), spikelet base (A2, A15, A17, A21), and floret (A11, A12) (Fig. S6B). Notably, cell types in snRNA-seq and snATAC-seq did not exhibit a straightforward one-to-one correspondence, reflecting the complexity of transcriptional regulation, likely including the role of histone modifications like H3K27me3 (Fig. S6C).

**Fig. 4.**
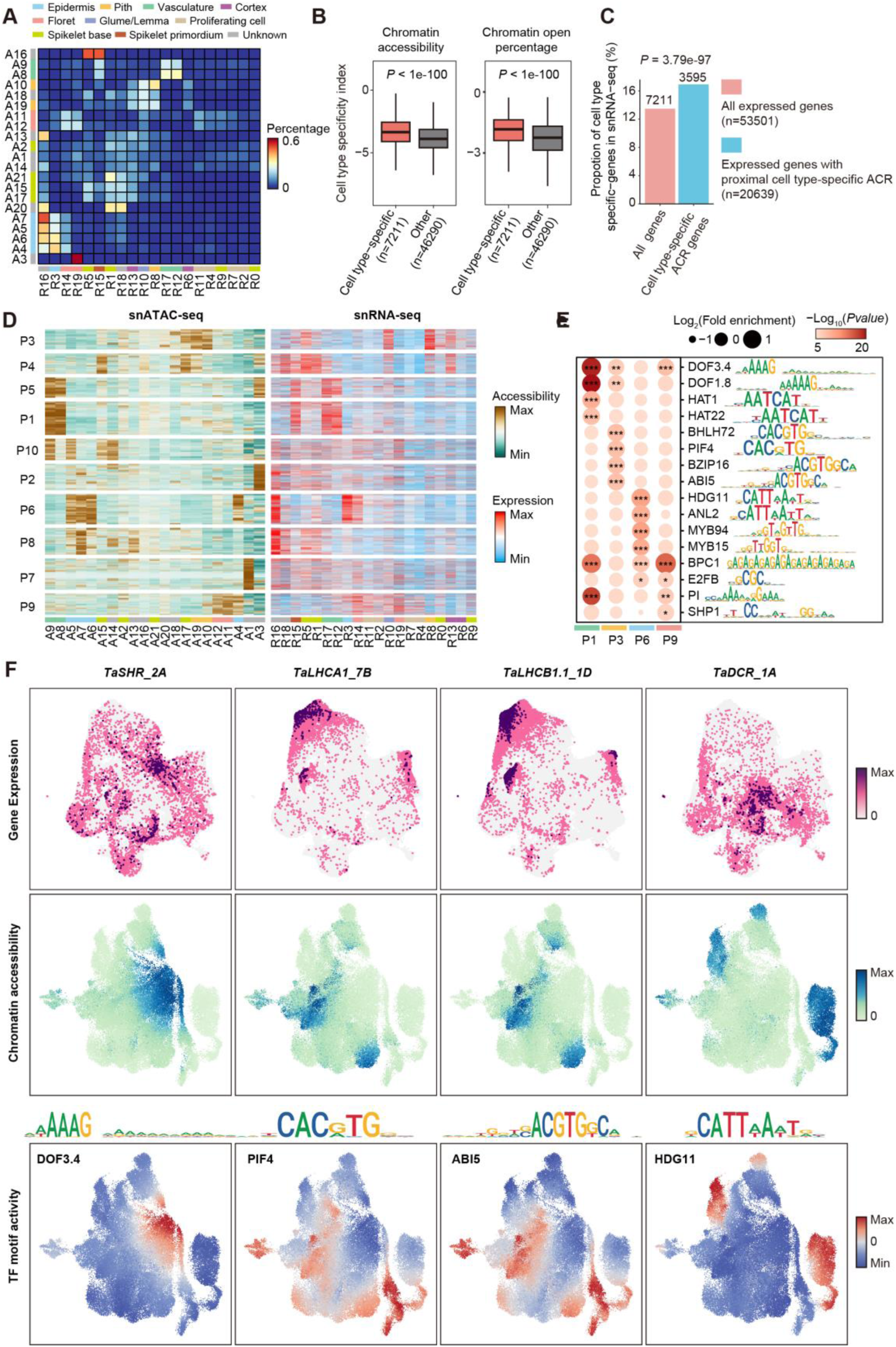
Cell-type specific transcriptional regulation during wheat inflorescence development. **A** Heatmap displaying the overlap proportion of cell type-specific genes in snATAC-seq with cell type-specific genes in different snRNA-seq cell types. Cell type annotations are shown on the left and bottom. **B** Cell type specificity index defined by chromatin accessibility values (left) and the proportion of cells where its chromatin is open (right) (two-sided Wilcoxon test). **C** Proportion of snRNA-seq cell type-specific genes among all expressed genes and genes whose proximal region has cell type-specific ACRs (two-sided Fisher’s exact test). Proximal ACR refers to ACRs located on gene promoters or genebodies. **D** Heatmap showing chromatin accessibility of cell type-specific ACRs in different cell types of snATAC-seq data, as well as the expression of proximal genes in various cell types of snRNA-seq. Cell type annotations are shown at the bottom and the color code are shown in panel A. **E** TF binding motif enrichment results in ACRs of different clusters in panel e (two-sided Fisher’s exact test, *P* < 0.05). The corresponding cell type annotations are shown at the bottom and the color code are shown in panel A. **F** Profiles of gene expression and chromatin accessibility of gene *TaSHR_2A*, *TaLHCA1_7B*, *TaLHCB1.1_1D*, *TaDCR_1A* and motif activity of TFs binding to csACRs on their promoters.

To assess gene expression regulation by chromatin accessibility at cellular level, we quantified gene expression and chromatin accessibility across cell types (Table S14, S15). A cell type specificity index was used to measure the specificity of expression and chromatin accessibility, with higher values indicating stronger cell type-specific patterns (Table S16). Genes with higher specificity indices for expression also tended to show greater specificity in chromatin accessibility (Fig. 4B, Fig. S6D). Notably, cell type-specific genes identified by snRNA-seq accounted for 13% of all expressed genes (Fig. 4C). However, this proportion increased to 17% among genes with cell type-specific ACRs in their proximal regions (*P* < 0.001, two-sided Fisher’s exact test), highlighting the significant role of these ACRs in regulating gene expression specificity (Fig. 4C).

Focusing on 3,595 genes with proximal cell type-specific ACRs, clustering analysis identified 10 distinct clusters (P1-P10) based on chromatin accessibility (Fig. 4D, Table S17). High chromatin accessibility in specific snATAC-seq cell types promoted the upregulation of corresponding genes. For example, cluster P3 contained ACRs predominantly open in pith cells (A10 and A19), located in the promoters of *Light-harvesting protein complex I* (*TaLHCA1*) and *TaLHCB1.1*, which are specifically expressed in pith (R8) (Fig. 4D, F, Fig. S6E). Genes specifically expressed in epidermis (R3) are regulated by ACRs opened in A4, A5, A6, and A7, such as those located in gene *TaDCR* and *TaFDH*’s promoter (Fig. 4D, F, Fig. S6E). Enrichment analysis of TF binding motifs within these ACRs revealed distinct regulatory patterns (Table S18). For instance, ACRs specific to vasculature cells (A8 and A9) in cluster P1 were enriched with motifs for DOF family TFs, and the motif activity of these TFs was also specifically elevated in vasculature (Fig. 4D-F). Floret-specific ACRs in cluster P9 were enriched with motifs for MADS and E2F family TFs, aligning with their roles in floret development (Fig.4D-F). These findings emphasize the role of cell type-specific ACRs and associated TFs in regulating spatiotemporal gene expression during wheat inflorescence development.

### cTRNs highlight node genes in shaping inflorescence architecture

To further understand cell fate determination during spikelet and floret formation, we integrated time-serial snRNA-seq and snATAC-seq data to construct cTRNs (Fig. 5A, Table S19, S20). The cTRNs revealed that TFs expressed in specific cell types not only regulate genes within their own cell type but also influences genes in neighboring cell types (Fig. 5A). For instance, TFs from the E2F and CPP families, highly expressed in proliferating cells, not only regulate cell cycle related genes but also regulate genes such as *VRT2*, *TaTFL1*, and *TaSEP3*, which are abundant in R5 (spikelet base), R15 (spikelet primordium), and R14 (floret) (Fig. 5A). These TFs may maintain cell identity while facilitating cell fate transitions. We identified common TF families, including bZIP, HD-ZIP, and MADS-box, as well as specific families like AP2 and SBP, within the cTRNs for spikelet and floret formation, respectively (Fig. 5A). Notably, during spikelet formation, factors related to auxin and cytokinin signaling played prominent roles, such as auxin influx carriers LIKE AUXIN RESISTANT 1 (TaLAX1), efflux carriers TaPIN1, signaling regulators indole-3-acetic acid inducible 27 (TaIAA27), and cytokinin synthesis enzymes LONELY GUY 6 (TaLOG6) and TaLOG8 (Fig. 6A). In contrast, the cTRNs for floret formation were dominated by MADS-family TFs, including TaAGL6, TaSEP3, APETALA 3 (TaAP3), TaPI, and TaAG, underscoring their central role in floret fate determination (Fig. 5A). These findings highlight the critical roles of hormones in inflorescence patterning and various TFs in spikelet and floret differentiation.

**Fig. 5.**
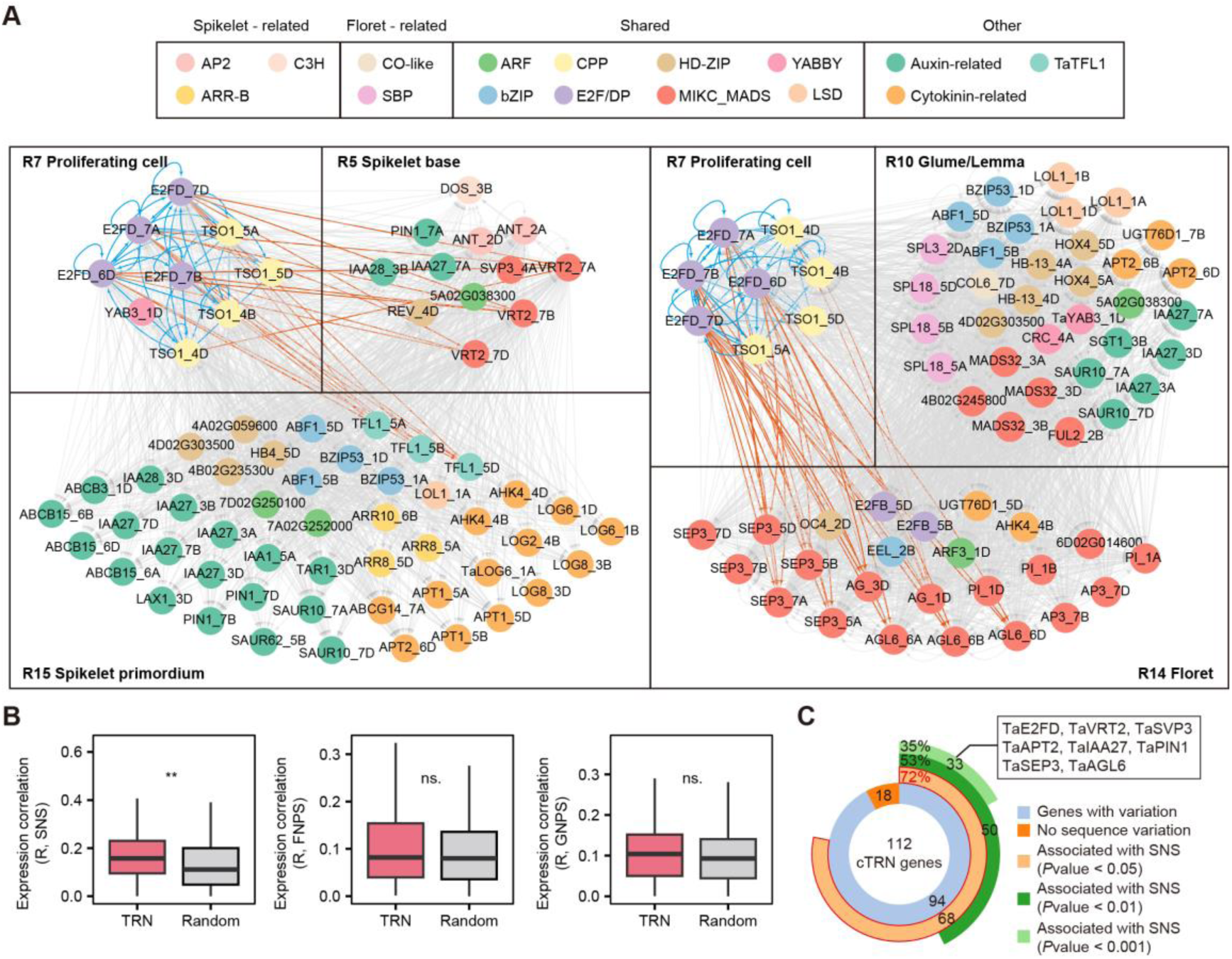
cTRN underlying wheat spikelet and floret formation. **A** Cell type-specific TRN underlying wheat spikelet and floret formation. The red lines represent the regulation of genes with high expression in R7 by ERF and CPP family, while the blue lines represent their regulation of genes with high expression in other cell types. **B** The association between gene expression level and SNS, FNPS, and GNPS (two-sided Wilcoxon test). TRN represents TRN node genes. Random refers to genes selected randomly. **C** The association between sequence variations in TRN genes and SNS.

**Fig. 6.**
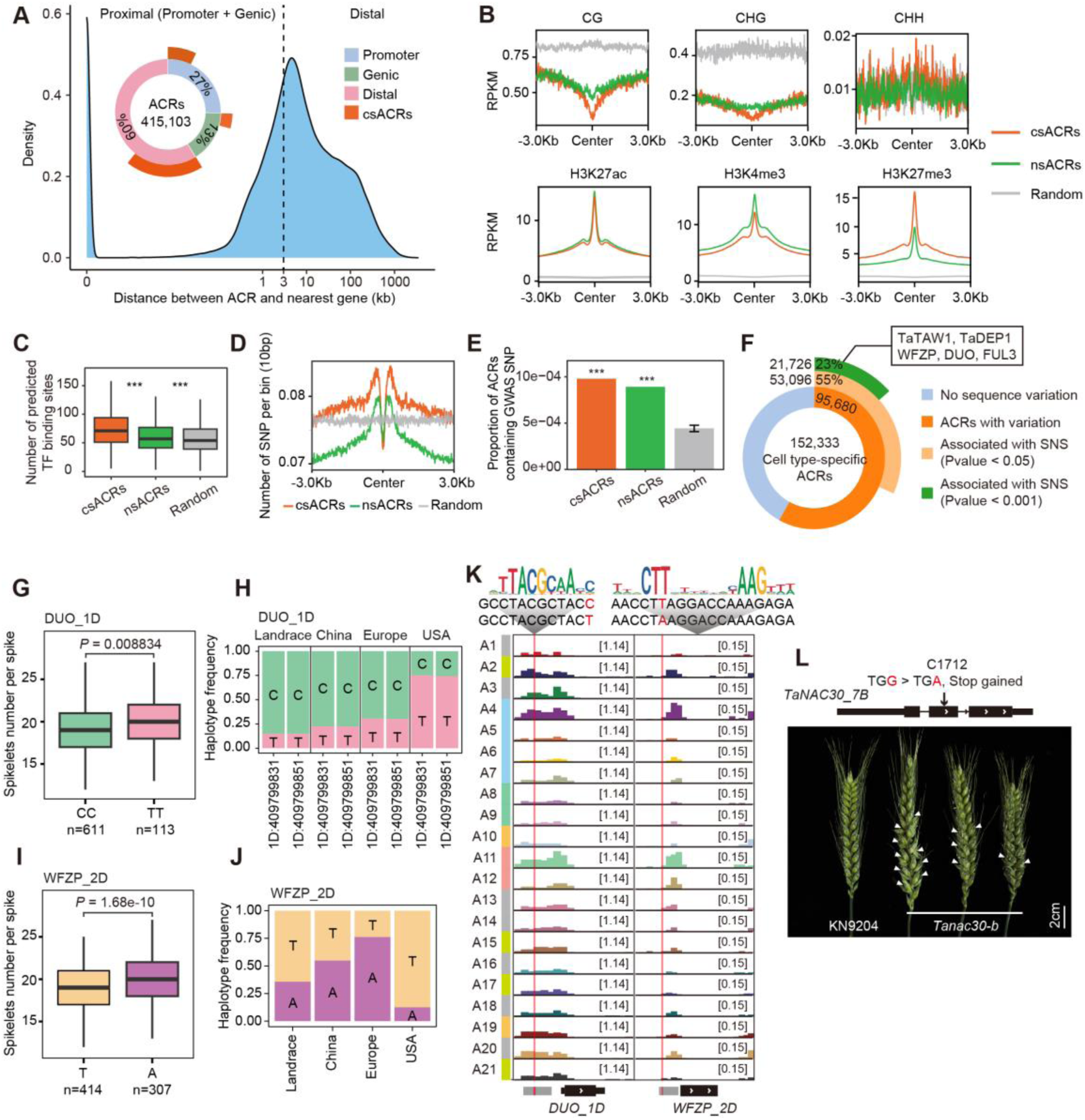
Cell type specific ACRs for regulating spike architecture. **A** Distribution and proportion of ACRs in different genomic locations. Promoter refers to the region from 3,000 bp upstream of transcription start site (TSS) to TSS. Genic refers to the region from TSS to transcription end site (TES). The other regions are defined as Distal. **B** Distribution of DNA methylation and histone modification across cell type-specific ACRs, non-specific ACRs and random regions. Random represents 300,000 sequences of 500 bp length randomly selected form regions outside the gene coding region and ACR. **C** Number of predicted TF binding sites detected within ACRs. **D** Distribution of the number of SNPs across ACRs. **E** Proportion of ACRs that overlap with reported spike-related GWAS SNPs. **F** Association between sequence variations within cell type-specific ACRs and SNS. **G, I** Spikelet number per spike of different haplotypes divided by SNPs within csACRs located in the promoter of *DUO_1D* (**G**) and *WFZP_2D* (**I**) (two-sided Student’s t-test). **H, J** Haplotype frequency of SNPs within csACRs located in the promoter of *DUO_1D* (**H**), and *WFZP_2D* (**J**). **K** Chromatin accessibility profiles of *DUO_1D* and *WFZP_2D* across snATAC-seq cell types. Cell type annotations are shown on the left and the color code are shown in Fig. 3a. Red box shows the location of NAC motif. **L** Photograph comparing spike of KN9204 and *Tanac30-b* mutant.

Recognizing the central role of node genes in TRNs (Davidson et al. 2002; Lin et al. 2024), we examined the association of 112 node genes within these cTRNs with spike-related traits. Analysis of population transcriptomes from the double ridge stage (Wang et al. 2017) revealed that node genes in the cTRNs correlated more strongly with SNS than random genes, but showed no significant associations with FNPS or GNPS (Fig. 5B). Notably, 68 of 94 node genes with sequence variations were associated with SNS, including significant hits (*P* < 0.05) for known regulators of inflorescence architecture such as *VRT2*, *TaSVP3*, and *TaAGL6*, as well as new candidates like *TaE2FD*, *TaPIN1*, *ADENINE PHOSPHORIBOSYL TRANSFERASE 2* (*TaAPT2*), and *TaIAA27* (Fig. 5C, Table S21). These findings underscore the direct involvement of node genes within cTRNs in regulating spikelet number and wheat inflorescence development.

### Importance of cell type-specific ACRs in shaping wheat inflorescence architecture

In addition to *trans*-factors, *cis*-regulatory elements play a crucial role in spatiotemporal gene expression. Using snATAC-seq data, we identified 415,103 ACRs (Table S22), with 152,333 being cell type-specific ACRs (csACRs) (Table S23), predominantly located in intergenic regions (Fig. 6A). These csACRs exhibited lower CG and CHG DNA methylation (Fig. 6B) but higher TF binding density (Fig. 6C) compared to non-specific ACRs (nsACRs), highlighting their role in modulating gene expression. Active histone marks H3K27ac and H3K4me3 showed similar levels between csACRs and nsACRs, while repressive histone mark H3K27me3 was higher in csACRs (Fig. 6B), consistent with its role in cell fate determination (Xiao and Wagner 2015).

To investigate the association between csACRs and spike-related traits, we utilized resequencing data from 1,051 Watkins landrace collections and modern cultivars (Cheng et al. 2024), along with genome-wide association (GWAS) results from previous studies on traits such as spike length, SNS, and GNPS (Table S24). The sequences of ACR were more conserved than random sequences (Fig. 6D) but contained a higher number of SNPs linked to spike-related traits (Fig. 6E). Notably, csACRs harbored a greater proportion of SNPs associated with these traits (Fig. 6E), highlighting their potential role in trait regulation. Analysis of the Watkins population’s genotypic and phenotypic data revealed that 23% of csACRs contained variants (21,726) were significantly associated with SNS (*P* < 0.001) (Table S25). Genes like *TAWAWA1* (*TaTAW1*), *WFZP*, *DUO*, and *DENSE AND ERECT PANICLE 1* (*TaDEP1*), with orthologs known to regulate inflorescence architecture in rice (Huang et al. 2009; Taguchi-Shiobara et al. 2011; Yoshida et al. 2013; Du et al. 2021; Wang et al. 2022), contained SNPs in their csACRs that were significantly associated with SNS in wheat (Fig. 6F).

We further investigated whether these csACR variants have been selected during wheat breeding using public resequencing datasets (Niu et al. 2023; Cheng et al. 2024). Using cross-population composite likelihood ratio (XP-CLR) tests (Chen et al. 2010) with the top 5% as threshold, we identified 3,629, 3,706, and 3,485 potential selective regions in the breeding processes of China, Europe, and the USA, respectively. Notably, distinct selection patterns were observed for the *DUO* and *WFZP* genes (Fig. S7A, B), both associated with the supernumerary spikelets (Du et al. 2021; Li et al. 2021b; Wang et al. 2022). The haplotype linked to increased SNS in *DUO_1D* (TT) remained nearly neutral during wheat breeding in China and Europe but increased in frequency from 15% to 75% in the USA (Fig. 6G, H). In contrast, the haplotype linked to increased SNS in *WFZP_2D* (A) was positively selected in China and Europe, with frequencies rising from 35% to 55% and 76%, respectively, but negatively selected in the USA (Fig. 6I, J). These findings suggest that wheat breeders in different regions may have selected csACRs that regulate distinct genes, yet have similar effects on wheat inflorescence structure, emphasizing the complex, region-specific nature of wheat breeding strategies.

The SNPs significantly associated with SNS in the csACRs of *DUO_1D* and *WFZP_2D* promoters also affect the binding of different NAC sub-family TFs (Fig. 6K). Additionally, we identified the EMS TILLING line C1712 with a premature mutation in *TaNAC30* (Wang et al. 2023), exhibiting supernumerary spikelets similar to DUO and WFZP mutants (Fig. 6M) (Poursarebani et al. 2015; Du et al. 2021; Li et al. 2021b; Wang et al. 2022). TaNAC30 positively regulates *WFZP* expression, and this regulation is diminished in the *Tanac30-b* mutant, which also displays the supernumerary spikelets phenotype. These findings highlight the crucial role of *cis-* and *trans-*regulatory mechanisms mediated by csACR in regulating wheat inflorescence architecture.

## Discussion

The inflorescence structure of wheat affects yield by influencing the number of spikelets and florets (Gauley and Boden 2019; Sakuma and Schnurbusch 2020). However, due to the complexity and asynchronous development of the wheat inflorescence, studying the formation of spikelets and florets has been challenging (Backhaus et al. 2022, 2023). In this study, we integrated multiple single-cell omics technologies to decode the cellular transcriptional regulatory networks underlying wheat inflorescence development with spatiotemporal resolution.

### Tackling wheat inflorescence development through spatiotemporal analysis

SnRNA-seq provides gene expression profiles at the cellular level, but its full potential is limited due to the loss of spatial information during library preparation (Adema et al. 2024). Spatial transcriptomics technologies, such as scStereo-seq and MERFISH, can compensate for this limitation and enable the investigation of spatiotemporal development in complex tissues (Xia et al. 2022; Long et al. 2024). Unlike the well-defined marker genes and complete cell lineages found in *Arabidopsis* roots (Benfey and Scheres 2000), wheat lacks clear marker genes, and the cell lineage during inflorescence development is still unclear. Moreover, the development of spikelets at different locations along the wheat inflorescence occurs asynchronously, which requires position information provided by spatial transcriptomics to annotate cell type and investigate the developmental asynchrony. While integrating spatial transcriptomics with snRNA-seq to investigate the development of complex tissues, such as the heart, brain, and intestine (Moffitt et al. 2018; Asp et al. 2019; Fawkner-Corbett et al. 2021), has become common in animals, studies integrating these two approaches in plants remain limited.

In this study, we tackled these challenges in wheat inflorescence development by leveraging both high-throughput single-cell spatial transcriptomics (scStereo-seq) and high-sensitivity MERFISH with snRNA-seq. ScStereo-seq provided spatial information on all cell types and gene expression, while MERFISH offered more detailed spatial gene expression data. For instance, based on the spatial expression specificity of *TaBHLH10* in MERFISH, we identified the ovary, a rare cell type at W5 stage (Fig. 2). By integrating these techniques, we constructed a comprehensive spatiotemporal gene expression atlas at the cellular level, and annotated 20 cell types within three major distinct groups with spatiotemporal context for wheat inflorescence development (Fig. 2). This atlas provides valuable data resources for studying inflorescence development in wheat and other crops.

### Developmental trajectory of spikelet and floret divergence

The architecture of plant inflorescences is largely shaped by the branching patterns of meristems before their terminal differentiation into florets (Zhu and Wagner 2020; Koppolu et al. 2022). The “transient model” of inflorescence development suggests that indeterminate meristems continuously generate lateral meristems until they acquire a determinate fate, influenced by vegetative (VEG) levels (Prusinkiewicz et al. 2007). This model explains the development of simple inflorescences, such as those in *Arabidopsis*. In Poaceae, the transient model interprets inflorescence branching as a process in which meristems acquire different identities in a programmed manner (e.g., wheat: IM-SM-FM; rice: IM-BM-SM-FM) (Kellogg 2022; Koppolu et al. 2022). These identities are thought to be controlled by homeotic genes (McSteen et al. 2000). However, the absence of homologous meristem identity genes in Poaceae and the finding that the *BRANCHED SILKLESS 1/ FRIZZYPANICLE* (*BD1/FZP*) gene, a candidate for SM identity, is not expressed in SMs, complicate this model (Chuck et al. 2002; Komatsu et al. 2003; Li et al. 2021b). Whipple proposed an alternative “signal center model”, suggesting that a signal center adjacent to the meristem, rather than the meristem itself, determines its fate (Whipple 2017).

In our single-cell analysis of wheat, we did not identify distinct “meristem” cells but focused on spikelet primordium (R15) and floret (R14) cells to examine inflorescence development. Our trajectory inference revealed no direct developmental trajectory from spikelet primordium to floret cells (Fig. 3). Instead, both originate from proliferating cells at the W3 and W5 stages, respectively. This finding contrasts with the “transient model” but aligns with observations in both wheat and rice. In wheat, Long et al. observed the coordinated transcription of spikelet ridge and leaf ridge, suggesting that leaf ridge may serve as a signaling center during meristem transition. In rice, Zong et al. identified distinct trajectories for IM-SM and IM-BM contrasting the IM-BM-SM sequence in the “transient model”. While our wheat data does not conclusively support the “signal center” model, it highlights the limitations of the “transient model” and suggests additional insights in Poaceae inflorescence development.

### Trans- and cis- regulation in shaping inflorescence structure and potential application in breeding process

Inflorescence development is not regulated by a single gene but is mediated by a complex regulatory network (Kellogg 2022). Within these networks, cis-regulatory elements and trans-acting factors coordinate to control the spatiotemporal expression specificity of target genes (Lin et al. 2024). Using time-series snRNA-seq and snATAC-seq data, we constructed the cTRNs for spikelet and floret formation, revealing 68 genes with sequence variations significantly associated with spike traits (Fig. 5). Additionally, in the csACRs that mediate the binding of trans-acting factors to target genes, 53,096 ACRs contain sequence variations significantly correlated with spike traits (Fig. 6). For example, SNPs significantly associated with spikelet number were found in the csACRs of the promoters of genes regulating multiple spikelet formation, such as *WFZP* and *DUO* (Fig. 6). These SNPs may regulate the multi-spikelet phenotype by affecting the binding of upstream NAC family transcription factors. These provide potential targets for the improvement of wheat inflorescence architecture, including both cis-elements and trans-acting factors.

While TFs typically regulate multiple target genes, CREs exhibit more specific regulation, making them more amenable to precise spatiotemporal control via CRISPR (Signor and Nuzhdin 2018). This is particularly important for pleiotropic genes, such as *IPA1* in rice and *WOX9* in tomato, which are expressed in multiple tissues or cell types (Hendelman et al. 2021; Song et al. 2022). Editing their CREs allows precise expression control in specific tissues or cell types, minimizing negative pleiotropic effects (Swinnen et al. 2016; Saeed et al. 2022). Therefore, editing these csACRs or targeting the cell type-specific genetic circuits between TFs and csACRs may enable more precise improvement of wheat inflorescence architecture, eventually enhancing the grain yield.

## Methods

### Plant materials and growth conditions

The winter wheat cultivar KN9204 was used as the material in this study. Germinated seeds were vernalized at 4 °C for 30 days. Then, the seedlings were transplanted into soil and grown in a greenhouse at 22 °C/20 °C day/night, under long day conditions (16 h light/8 h dark). The stage-specific shoot apex of wheat was dissected under a stereomicroscope based on anatomical and morphological features, immediately frozen in liquid nitrogen, and stored at -80 °C. About 5 to 20 spikes were pooled for each of replicate of snRNA-seq and snATAC-seq libraries at six development stages.

### Nucleus isolation

The nuclei isolation procedure was modified from the product information sheet of the CelLyticTM PN Isolation/Extraction Kit (Sigma). Briefly, plant tissues were chopped and mixed into 1× NIBTA buffer (1× NIB, 1 mM dithiothreitol, 1× ProtectRNATM RNase inhibitor, 1× cOmplete™, ethylenediaminetetraacetic acid-free Protease Inhibitor Cocktail, and 0.3% Triton X-100) on prechilled plates, and homogenates were transferred to 15 mL tubes. The tubes were then shaken for 5 minutes on ice, after which lysates were passed through 40 μm strainers, and flow-throughs were collected into new 15 mL tubes. These were then centrifuged for 10 minutes at 1,260 × g, the supernatants were decanted, and the pellets were resuspended in 4 mL of 1× NIBTA buffer. In new 15 mL tubes containing 80% Percoll® solution (4 mL Percoll® plus 1 mL NIBTA buffer), lysates were carefully overlaid onto the Percoll® layers. Tubes were centrifuged for 30 minutes at 650 × g, after which most of the nuclei had banded at the 1× NIBTA buffer and Percoll® interface. The nuclei bands were gently collected into new 15 mL tubes, 10 mL of 1× NIBTA buffer was added, and the tubes were recentrifuged for 5 minutes at 1260 × g. Nuclei pellets were then washed twice in 1× NIBTA buffer and resuspended in PBS containing 0.04% BSA (for snRNA-Seq) to a final concentration of 2,000 nuclei/uL or PBS containing 1% BSA (for snATAC-Seq) to a final concentration of 6,250 nuclei/uL.

### snRNA-seq library construction

The snRNA-seq libraries were prepared using the DNBelab C Series High-throughput Single-Cell RNA Library Preparation Kit (MGI, 940-000047-00) as previously described (Liu et al. 2019). Sequencing was performed on MGI 2000, DNBSEQ T7, and DIPSEQ T1 plantforms.

### snATAC-seq library construction

The snATAC-seq libraries were prepared using the DNBelab C Series Single-Cell ATAC Library Prep Set (MGI, #1000021878) as previously described (Lei et al. 2022). Sequencing was performed on MGI 2000, DNBSEQ T7, and DIPSEQ T1 platforms.

### scStereo-seq library construction

The scStereo-seq libraries were prepared as previously described (Chen et al. 2022). Briefly, the plant tissue was embedded in OCT and sectioned into 10-micrometer slices. These slices were then adhered to the Stereo-seq chip, which was incubated at 37°C for 3 minutes. Subsequently, the tissue was fixed in methanol at -20°C for 30 minutes. The cell walls were imaged using a moticfluorescence microscope, leveraging the intrinsic fluorescence properties of the plant tissue. The subsequent experimental steps, including permeabilization, reverse transcription, tissue removal and cDNA release, and library preparation, were carried out as previously described (Xia et al. 2022). Final sequencing was performed on the DIPSEQ T1 platform.

### snRNA-seq data analysis

Raw data of snRNA-seq was processed using an open-source pipeline (https://github.com/MGI-tech-bioinformatics/DNBelab_C_Series_HT_scRNA-analysis-software) to generate the read count matrix for each gene and cell. Firstly, raw sequencing reads were filtered to exclude those with an average base quality score below 4, more than two bases with a quality score below 10, reads containing N bases, or reads with invalid barcodes. The filtered reads were then demultiplexed based on barcode assignment. Subsequently, the filtered reads were aligned to the wheat reference genome (IWGSC RefSeq v1.0 ((International Wheat Genome Sequencing Consortium (IWGSC) 2018) using STAR (v2.7.9a) (Dobin et al. 2013) and annotated with the gene set from IWGSC Annotation v1.1 using PISA (Shi et al. 2022). Valid nuclei were identified using the "BarcodeRank" function of the DropletUtils (v1.16.0) (Lun et al. 2019), which filtered out background beads and beads with UMI counts below 500. Finally, PISA was employed to generate a UMI count matrix for each cell and construct a gene-by-cell matrix for each snRNA-seq library.

Downstream analyses were performed using Seurat (v4.3.0) (Hao et al. 2021). First, Seurat objects were created for each sample using the “CreatSeuratObject” function with the following filter criteria: min.cells=3, min.features=2,000. The individual Seurat objects were then merged into a single Seurat object, and data normalization was performed using “SCTransform” function. The top 3,000 variable features were selected for PCA using the “RunPCA” function. The first 30 dimensions were used for clustering analysis, which was conducted using the “FindNeighbors” and “FindClusters” functions with a resolution set to 0.5. Finally, the “RunUMAP” function was employed for visualization. Cell typs-specific genes for each cell type were identified using the “FindAllMarkers” function with parameters set to “only.pos=T, min.pct=0.1, logfc.threshold=0.25”.

### snATAC-seq data analysis

The raw reads were processed by PISA (Shi et al. 2022), and then aligned to the wheat reference genome (IWGSC RefSeqv1.0) using BWA (v 0.7.17-r1188) (Li 2013), resulting in BAM files. Fragment files for each snATAC-seq library were then generated using bap2 (Lareau et al. 2019). Downstream analyses were performed in ArchR (v1.0.2) (Granja et al. 2021). All samples were merged to create Arrow files using the “createArrowFiles” function, specifying the promoter region as 3,000 bp upstream to 500 bp downstream of the TSS. Cells were filtered based on the following criteria: minTSS=5, minFrag=3,000, maxFrags=1e+5. Doublets were identified and removed using the “doubScores” and “filterDoublets” functions with default parameters. Dimension reduction and clustering analyses were performed using the “addIterativeLSI” and “addClusters” functions, respectively, with a resolution set to 0.6. To visualize these results, the UMAP matrix is added to the ArchR project using the “addUMAP” function. Cells within the same cluster were grouped into pseudo-samples to identify ACRs using MACS2 (v2.2.7.1) (Zhang et al. 2008). Cell type-specific genes and cell type-specific ACRs for each cell type were identified using the “getMarkers” function with parameter “maxCells = 2000” and thresholds “FDR <= 0.05 & Log2FC >= 1”. The identified ACRs were annotated to the wheat genome using the R package ChIPseeker (v1.24.0) (Yu et al. 2015).

### scStereo-seq data analysis

Raw data of scStereo-seq were processed as previously described (Chen et al. 2022). Coordinate identities (CIDs) were assigned by mapping to the coordinates of the in situ captured chip, allowing for one base mismatch to account for potential sequencing or PCR errors. After removing reads with UMIs that had a quality score below 10, the remaining reads were aligned to the wheat reference genome (IWGSC RefSeq v1.0) and annotated to the gene set (IWGSC Annotation v1.1). An expression profile matrix containing CID information was then generated for subsequent analyses. A Seurat object was created by Seurat (v4.3.0) for each sample, and cells were filtered with the following criteria: min.cells = 3, min.features = 100.

Autofluorescence images of a section were used to identify the cell regions in scStereo-seq data generated from the same section. The process began with the generation of a grayscale map of the scStereo-seq data, where each pixel represented one DNB. A global threshold was applied to filter out background noise in the registered images, creating masks for segmentation. A Gaussian-weighted local threshold was then computed to indicate cell locations, using a block size of 41 and offset of 0.03. To address overlapping cells, an exact Euclidean distance transformation was performed, and local peaks were identified as markers with a minimum distance of 15 pixels. The Python package Cellpose (v2.1.0) (Pachitariu and Stringer 2022) was utilized to segment the autofluorescence images and generate cell-mask images. Manual registration between the maps and the cell-mask images was performed to align the pixels of the cell-mask images with DNB coordinates. Labels representing different cells were transferred to pinpoint DNBs corresponding to spatial positions by the watershed algorithm via scikit-image python package (v0.18.1) (Neubert and Protzel 2014). Although tissue damage resulted in the appearance of some false cells in the cell-mask images—specifically, large cells located at the damaged tissue areas—their proportion is minimal (less than 2%) (Fig. S2C). Therefore, in subsequent analyses, cells were filtered using UMI Count without removing these "false cells."

### Mapping snRNA-seq data onto scStereo-seq data

Tangram (v1.0.4) (Biancalani et al. 2021) was used to map the snRNA-seq data onto scStereo-seq data. A total of 2,867 top cell type-specific genes in snRNA-seq (p_val_adj < 0.05 & avg_log2FC > 0.5) were used as training genes. To obtain the spatial information of cell types in snRNA-seq, mapping was performed at the cluster level using the function “map_cells_to_space” with parameter mode = ‘clusters’. Subsequently, spatial probability maps of cell types were generated using the function “project_cell_annotations” with parameters: annotation= ‘seurat_clusters’ and threshold = 0.5. For each cell in scStereo-seq data, cell types from snRNA-seq data were assigned based on the highest spatial probability. To obtain the spatial expression of genes in snRNA-seq, mapping was performed at the cell level using the function “map_cells_to_space” with parmeter mode = ‘cell’, followed by the generation of mapped spatial gene expression data using the function “project_genes”. Spatial visualization of cell types and gene expression was performed using OpenCV (v4.10.0) (Bradski and Kaehler 2008).

### Multiplexed error-robust fluorescence *in situ* hybridization (MERFISH)

We used the raw data and segmentation outputs from W5 stage inflorescence from Long et al. 2024 and visualised selected genes using the MERSCOPE Visualizer Tool.

### Trajectory inference and pseudotime analysis

Trajectory inference was performed using Monocle3 (v1.3.1) (Qiu et al. 2017). The input data for Monocle3 was derived from the Seurat object. Partitioning analysis was conducted on all cells, starting with data preprocessing and normalization using the “preprocess_cds” function, followed by dimension reduction and clustering using the “reduce_dimension” and “cluster_cells” functions to obtain the cell partition results. For trajectory inference and pseudotime analysis, we used cells from R7.P4, R5, R15 for spikelet formation and cells from R7.P1, R10, R14 for floret formation. The data were first processed sequentially using functions “preprocess_cds”, “reduce_dimension”, and “cluster_cells”. Pseudotime analysis was then performed using the “learn_graph” and “order_cells” functions. The “graph_test” function was used to identify genes whose expression changed with pseudotime. Z-scaled expression values of genes were extracted and then clustered in R.

### Cellular TRN construction

The cTRNs for spikelet and floret formation were constructed following a modified approach based on previously reported methods (Liu et al. 2023).

Step1: Candidate gene selection

Trajectory inference and pseudotime analysis were first performed using cells belong to R7, R5, and R15 in partition P4 (for spikelet formation) or R7, R10 and R14 in partition P1 (for floret formation). Genes whose expression change with pseudotime (Moran’s I, qvalue < 1e-10) in cell type-specific genes of R5, R7, R10, R14, and R15 were selected as candidates.

Step2: Linking motifs to target genes

The sequence of ACRs located in the promoter or genebody of candidate genes were extracted, and motifs were scanned using fimo (5.2.0) (Grant et al. 2011). Motifs files for Arabidopsis were downloaded from the PlantTFDB (http://planttfdb.gao-lab.org/) (Jin et al. 2017).

Step3: Linking TFs to motifs

Given the motif data were derived from Arabidopsis, we established links between Arabidopsis TFs and wheat TFs using the following criteria: (1) TFs in both species belong to the same family, and (2) the e-value in BLASTP (v2.10.1) should be less than 1e-5, with sequence identity greater than 30%. This process allowed us to establish connections between TFs and their corresponding motifs.

Step4: Linking TFs to target genes

Using results from Step2 and Step3, we created direct links between TFs and their target genes, forming the foundation of cTRNs underlying spikelet and floret formation.

Step5: cTRN of TFs in enriched TF families and functional related genes

The cTRNs were refined to include only TFs from families that were enriched in different clusters (two-sided Fisher’s exact test, pvalue < 0.01). Additionally, genes involved in auxin and cytokinin, as well as known regulators of wheat inflorescence development were incorporated into the network.

### Association analysis between SNPs and SNS

Sequence variation and phenotypic data from 1,051 Watkins landraces and modern cultivars (Cheng et al. 2024) were used to analyze the association between DNA sequence variation within TRN node genes or csACRs and spikelet number per spike. First, DNA sequence variations located within TRN node genes or csACRs were extracted. The population was then stratified into subpopulations based on a single SNP. Finally, a two sided Student’s t-test was employed to compare the difference of spikelet number per spike between the two subpopulations.

### Identification of breeding selection regions

To identify potentially selected regions during wheat breeding in China, Europe, and the USA, two published resequencing datasets (Niu et al. 2023; Cheng et al. 2024) were merged by bcftools (v1.3.1) (Danecek et al. 2021), retaining DNA variants present in both datasets. These variants were then filtered by VCFtools (v0.1.16) (Danecek et al. 2011) with the parameters “--max-missing 0.1 --maf 0.01”. The composite likelihood approach (XP-CLR) (Chen et al. 2010) (https://github.com/hardingnj/xpclr) was applied to identify selected regions during breeding with the following parameters: --size 500000, --step 100000, --maxsnps 500. Landrace was used as the reference, with modern cultivars from China, Europe, and the USA as queries. Genomic regions were ranked based on XP-CLR scores, and those in the top 5% were considered to be under selection. Adjacent selected regions within 300kb were merged to define the final selected regions.

## Data availability

The raw sequence data reported in this paper have been deposited in the Genome Sequence Archive of the National Genomics Data Center (Chen et al. 2021; CNCB-NGDC Members and Partners 2022), China National Center of Bioinformation/Beijing Institute of Genomics, Chinese Academy of Science (CRA018137) and are publicly accessible at https://ngdc.cncb.ac.cn/gsa (https://ngdc.cncb.ac.cn/gsa/s/B3z0EsBX).

## Code availability

Code used for all processing and analysis is available in GitHub (https://github.com/xmliu01/Decoding-the-cTRN-govering-wheat-inflorescence-development-with-spatiotemporal-resolution)

## Acknowledgements

This research is supported by the National Key Research and Development Program of China (2021YFD1201500), Beijing Natural Science Foundation Outstanding Youth Project (JQ23026), the National Natural Science Foundation of China (31921005), the European Research Council (ERC-2019-COG-866328), the UK Biotechnology and Biological Sciences Research Council (BBSRC) Delivering Sustainable Wheat (BB/X011003/1) Institute Strategic Program, and the Gatsby Charitable Foundation.

## Author contributions

J.X. designed and supervised the research, J.X. and C.U. analyzed and refined the manuscript. X.M.L. performed data analysis, prepared figures and wrote the draft of manuscript. X.L.L. was responsible for sample collection, snRNA-seq, snATAC-seq and scStereo-seq libraries preparation. X.L.L. and J.M.K were responsible for snRNA-seq, snATAC-seq, scStereo-seq library construction and sequencing. K.A.L. and C.U. conducted MERFISH data analysis. J.J.Y. conducted the Luc reporter assays. C.C. was involved in omics data analysis. D.Z.W. contributed to population-level data analysis. All authors discussed the results and commented on the manuscript.

## Supplemental Figures

**Fig. S1.**
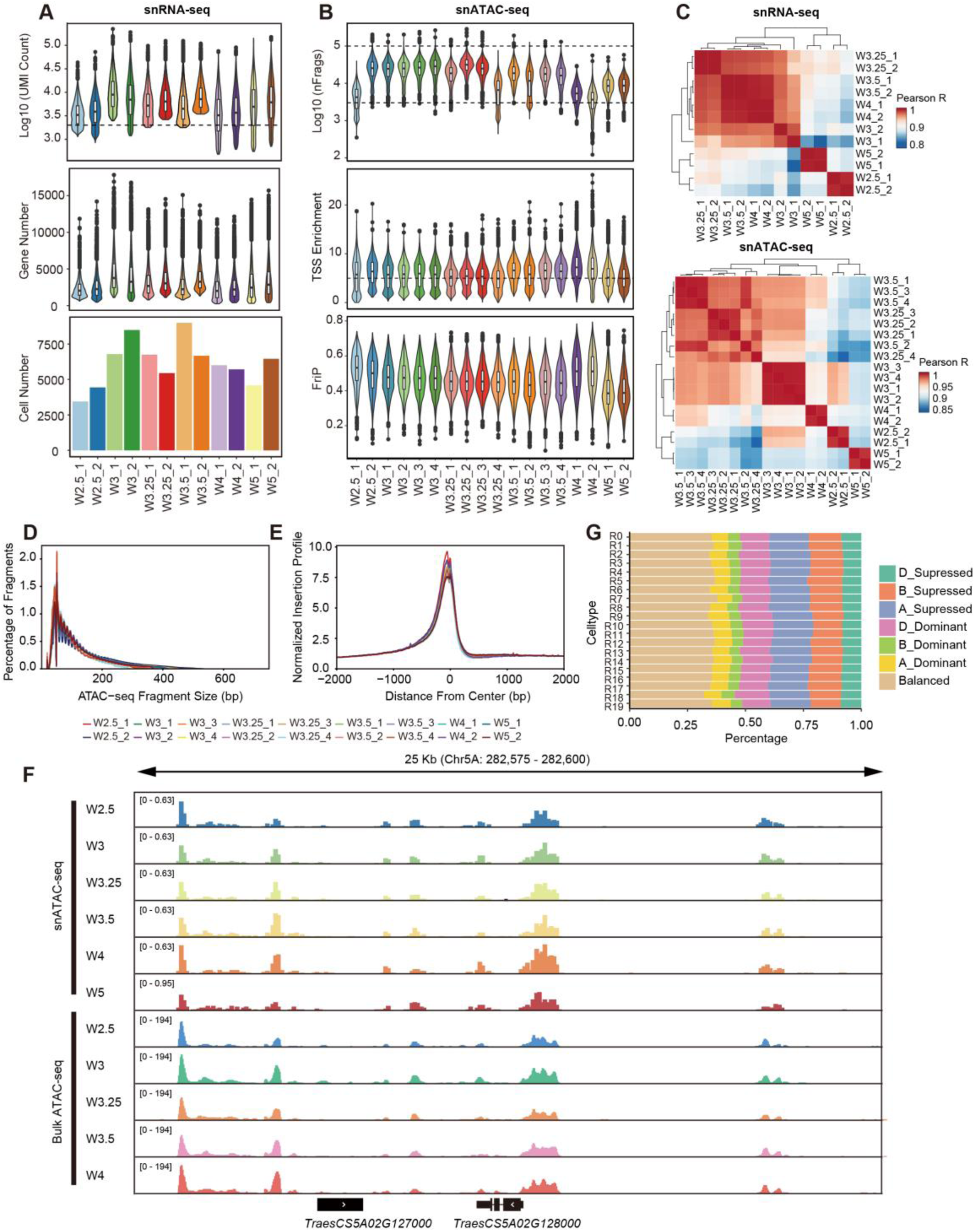
Data quality of snRNA-seq and snATAC-seq. **A** snRNA-seq quality metrics showing the distribution of UMI count, expressed gene number, and cell number in each library. The dashed line represents the filtering criteria. **B** snATAC-seq quality metrics showing the distribution of UMI count, TSS enrichment, and FriP in each library. The dashed line represents the filtering criteria. **C** Heatmap showing the Pearson correlation of each snRNA-seq and snATAC-seq library. **D** Fragment size distributions for each snATAC-seq library. **E** Normalized fragment count around transcriptional start sites for each snATAC-seq library. **F** Proportion of triads in each category of homoeolog expression bias across snRNA-seq cell types. “A_Dominant” represents the expression level of genes in the A subgenome being higher than their homoeologs in the B and D subgenomes; “A_supressed” represents the expression level of genes in the A subgenome being lower than their homoeologs in the B and D subgenomes; “Balance” represent the homoeologs from A, B, and D subgenomes being expressed at similar levels. **G** Chromatin accessibility profiles during wheat inflorescence development in bulk ATAC-seq and snATAC-seq data.

**Fig. S2.**
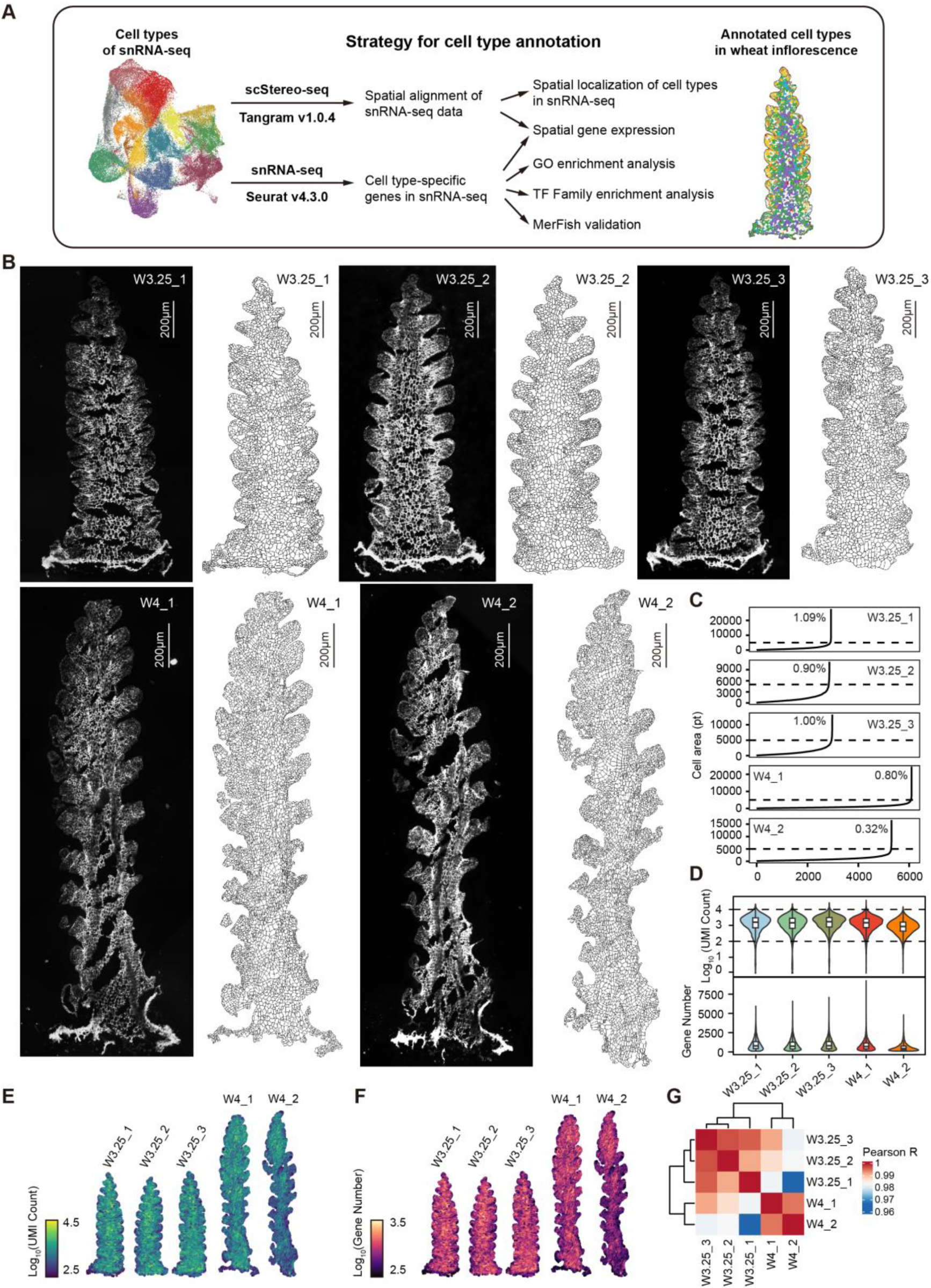
Data quality of scStereo-seq. **A** A diagram illustrating the strategy for cell type annotation in wheat inflorescence. **B** Cell wall staining images for each sample in scStereo-seq. **C** Cell area distribution across samples, with the dashed line representing 5,000 pt. The percentage indicates the proportion of cells larger than 5,000 pt. **D** The distribution of UMI count (**c**) and gene number (**d**) of each sample of scStereo-seq. **E, F** Spatial distribution of UMI count (**e**) and gene number (**f**) of each sample of scStereo-seq. **G** Heatmap showing the Pearson correlation of each scStereo-seq sample.

**Fig. S3.**
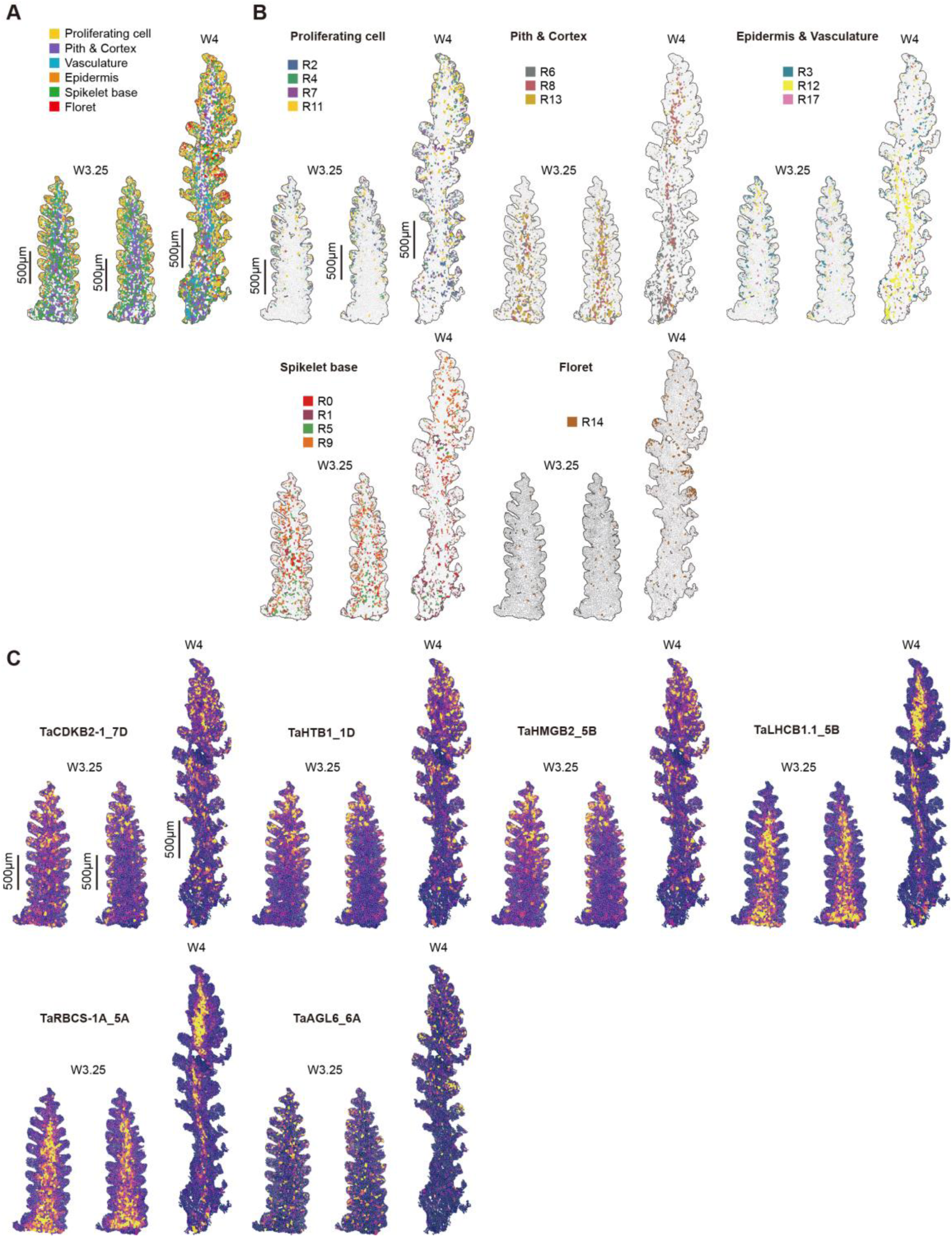
Spatial distribution of snRNA-seq data mapped on scStereo-seq data A, **B** Spatial distribution of cell types defined by snRNA-seq mapped on scStereo-seq data. **C** Predicted spatial gene expression of cell type-specific genes by mapping the snRNA-seq data to scStereo-seq data.

**Fig. S4.**
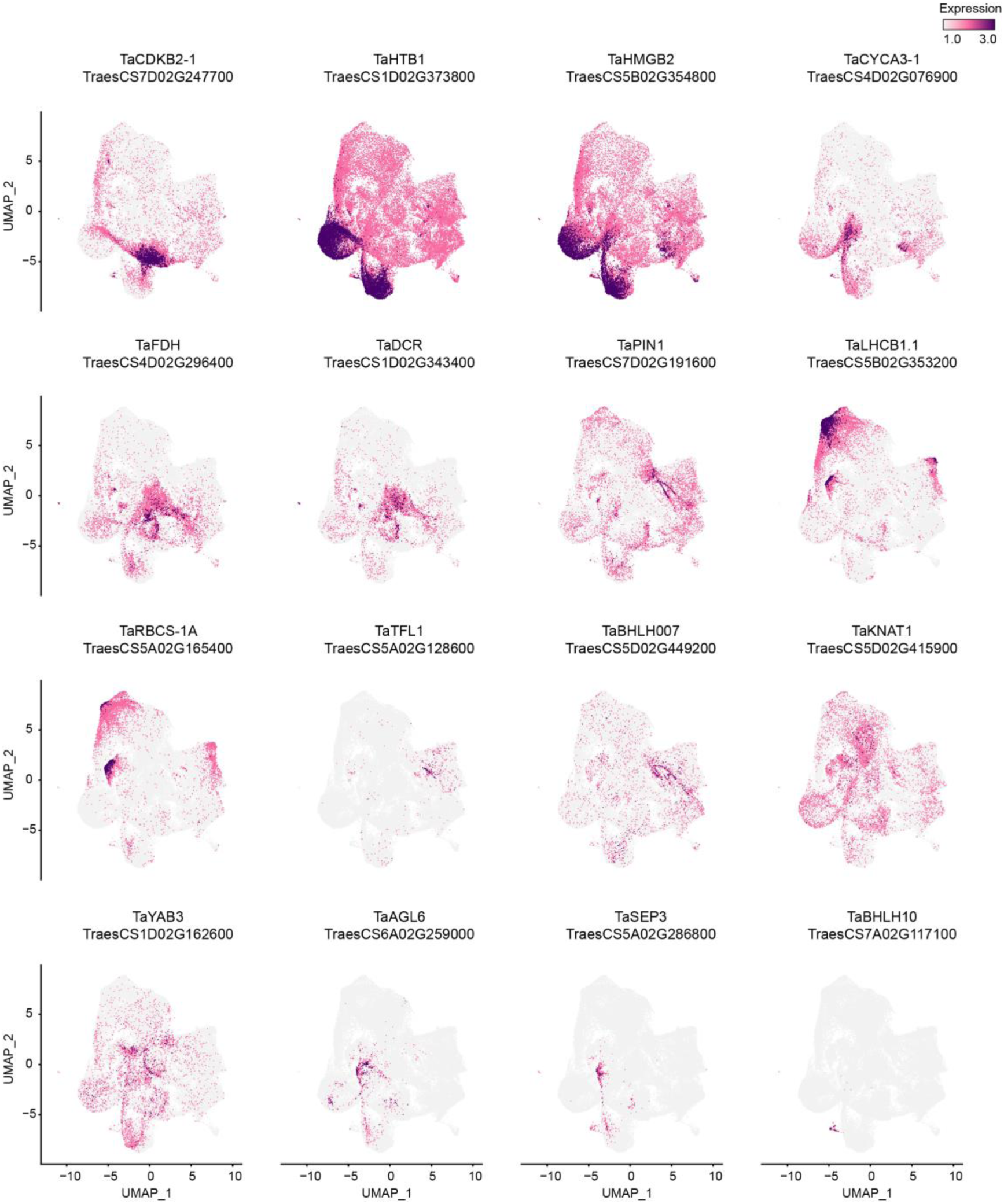
**Expression profiles of cell type-specific genes in snRNA-seq**

**Fig. S5.**
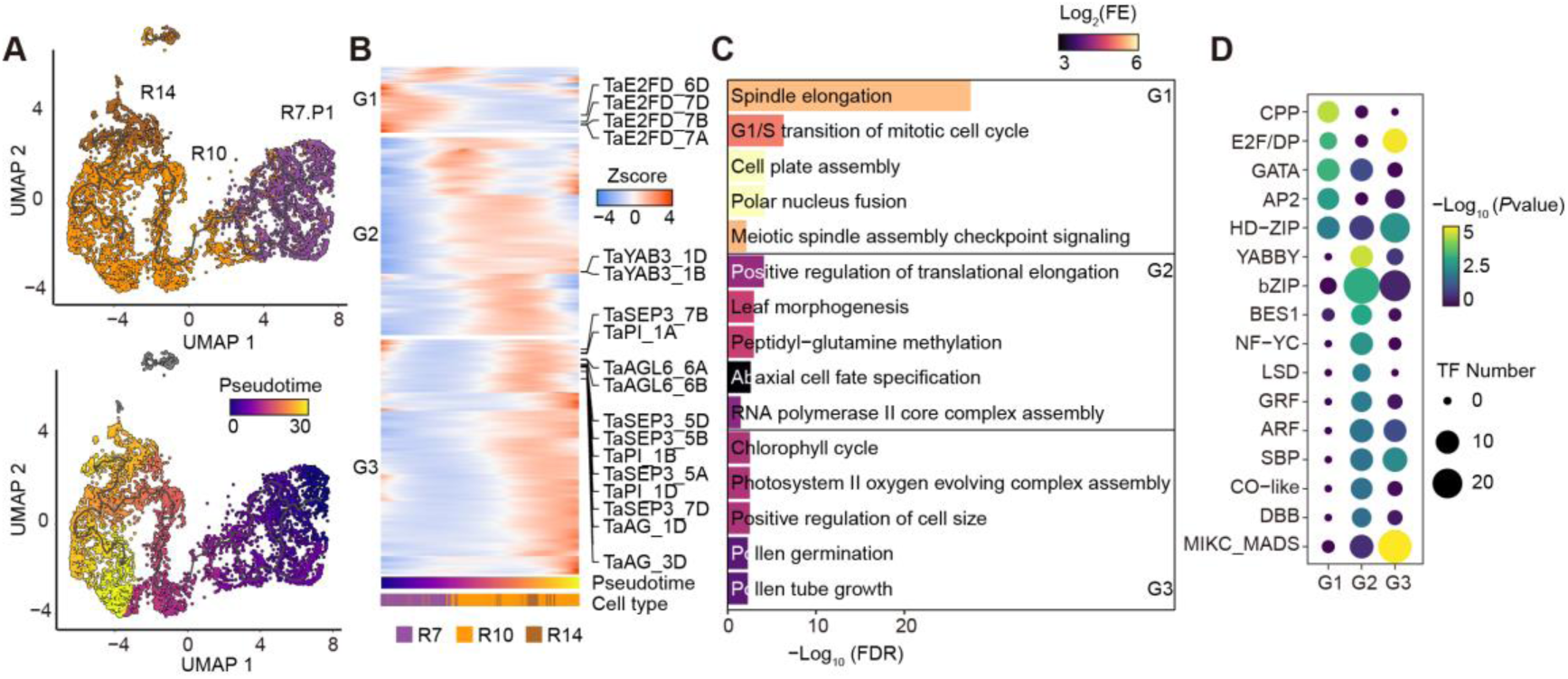
Trajectory inference for wheat floret formation. **A** Trajectory inference and pseudotime analysis of cells belonging to R7, R10, and R14 in partition1. **B** Heatmap showing the clustering results of genes whose expression change across pseudotime of floret formation. **C** GO enrichment analysis of genes in different clusters during floret formation (two-sided Fisher’s exact test, FDR for multiple testing, FDR < 0.05). FE: fold enrichment. **D** TF family enrichment of genes in different clusters during floret formation (two-sided Fisher’s exact test, *P*value < 0.01).

**Fig. S6.**
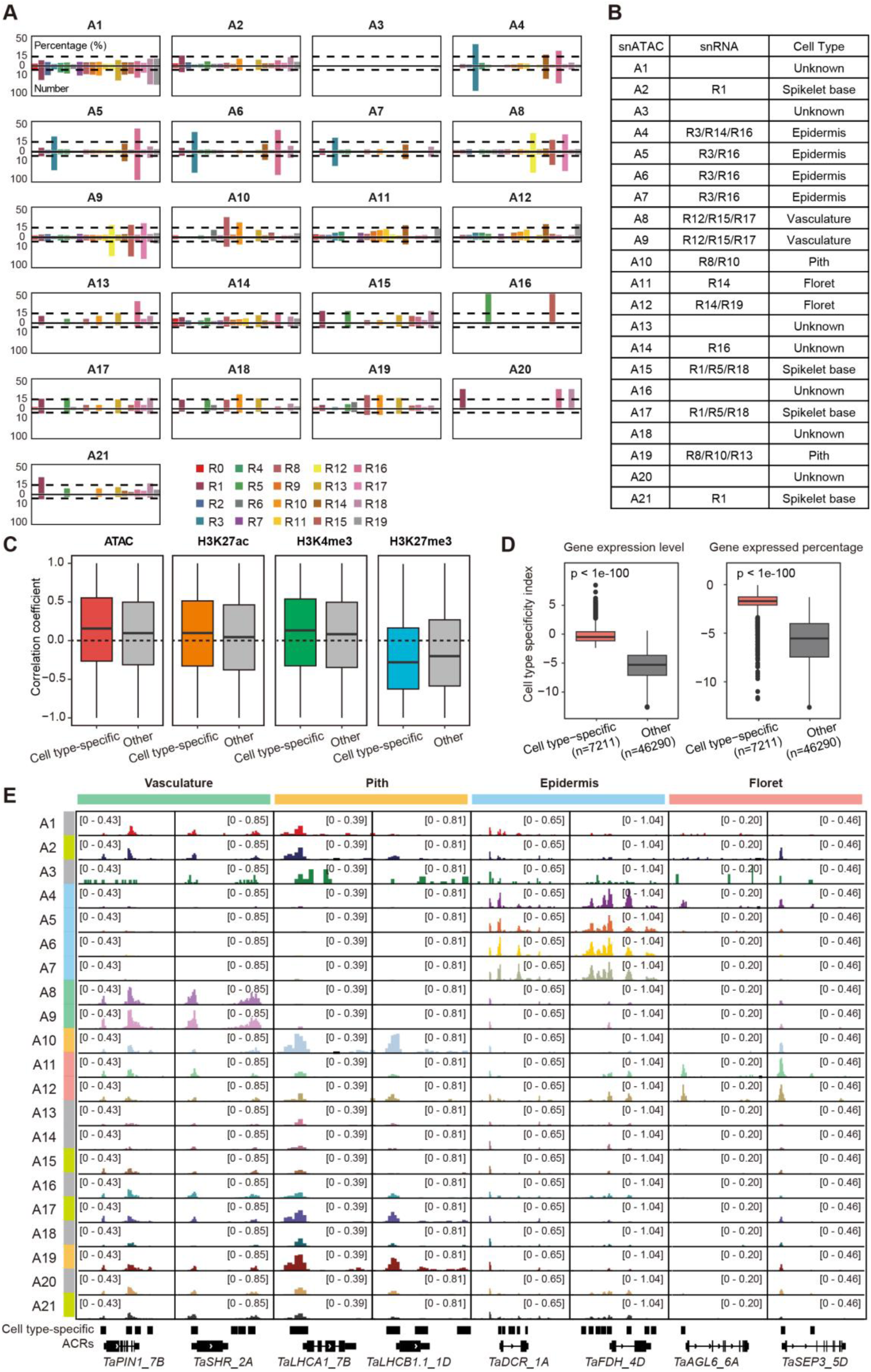
Correspondence between snRNA-seq and snATAC-seq. **A** Proportion of cell type-specific genes in snATAC-seq among the cell type-specific genes of each cell type in snRNA-seq. The dashed line represents the thresholds, with gene proportion no less than 15% and gene count no less than 10. **B** Correspondence between snATAC-seq and snRNA-seq cell types. **C** Pearson correlation coefficient between gene expression and changes in chromatin accessibility, histone modification of snRNA-seq cell type-specific genes, and other genes. Data on gene expression, chromatin accessibility, and histone modification are from published bulk data of wheat spike (Lin et al. 2024). **D** Cell type specificity index defined by gene expression values (left) and the proportion of cells expressing it (right) (two-sided Wilcoxon test). **E** Chromatin accessibility profiles of several cell type-specific genes across snATAC-seq cell types. Cell type annotations are shown on the left and the color code are shown in panel A.

**Fig. S7.**
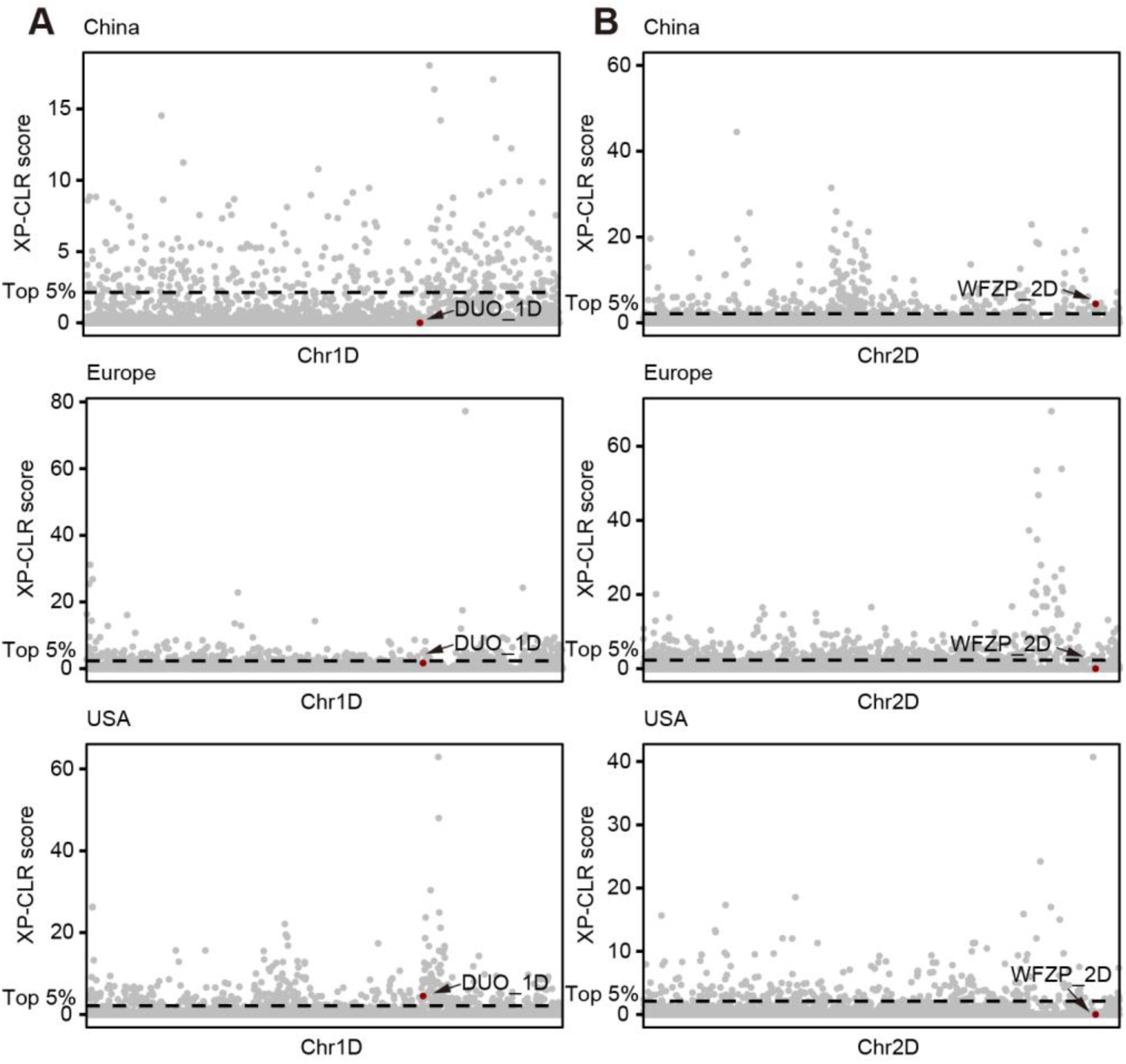
Profiling of selective regions during modern wheat breeding. **A, B** The selective regions during wheat breeding processes of China, Europe, and USA on the 1D chromosome (**A**), and 2D chromosome (**B**). The dashed lines indicate the cutoffs of the top 5% of XP-CLR score.

## Supplementary tables

**Table S1** Metadata of cells in snRNA-seq.

**Table S2** Metadata of cells in snATAC-seq.

**Table S3** Cell type-specific genes identified in snRNA-seq.

**Table S4** Cell type-specific genes identified in snATAC-seq.

**Table S5** Metadata of cells in scStereo-seq.

**Table S6** TF family enrichment results of cell type-specific genes in snRNA-seq.

**Table S7** GO enrichment results of cell type-specific genes in snRNA-seq.

**Table S8** Partition results of cells in snRNA-seq.

**Table S9** Differentially expressed genes between R7.P4 and R7.P1.

**Table S10** Pseudotime analysis results of cells belonging to R7, R5, and R15 in partition4.

**Table S11** Pseudotime analysis results of cells belonging to R7, R10, and R14 in partition1.

**Table S12** Clustering analysis results of genes whose expression change across pseudotime of spikelet formation.

**Table S13** Clustering analysis results of genes whose expression change across pseudotime of floret formation.

**Table S14** Gene expression levels and proportion of cells expressing each gene in snRNA-seq.

**Table S15** Chromatin accessibility of genes and proportion of cells with open chromatin in snATAC-seq.

**Table S16** The cell type specificity index of all expressed genes.

**Table S17** Clustering analysis results of proximal cell type-specific ACRs identified in snATAC-seq.

**Table S18** Enrichment of TF binding motifs within cell types-specific ACRs.

**Table S19** Cell type-specific TRN for spikelet formation.

**Table S20** Cell type-specific TRN for floret formation.

**Table S21** Association analysis between sequence variations within cTRN genes and SNS.

**Table S22** Annotation of ACRs identified in snATAC-seq.

**Table S23** Cell type-specific ACRs identified in snATAC-seq.

**Table S24** Reported GWAS loci associated with wheat spike-related traits.

**Table S25** Association analysis between sequence variations within cell type-specific ACRs and SNS.

## References

Adamski NM, Simmonds J, Brinton JF, Backhaus AE, Chen Y, Smedley M, Hayta S, Florio T, Crane P, Scott P, et al. Ectopic expression of Triticum polonicum VRT-A2 underlies elongated glumes and grains in hexaploid wheat in a dosage-dependent manner. Plant Cell. 2021:33(7):2296–2319. 10.1093/plcell/koab119

Adema K, Schon MA, Nodine MD, and Kohlen W. Lost in space: what single-cell RNA sequencing cannot tell you. Trends in Plant Science. 2024. 10.1016/j.tplants.2024.03.010

Ashikari M, Sakakibara H, Lin S, Yamamoto T, Takashi T, Nishimura A, Angeles ER, Qian Q, Kitano H, and Matsuoka M. Cytokinin Oxidase Regulates Rice Grain Production. Science. 2005:309(5735):741–745. 10.1126/science.1113373

Asp M, Giacomello S, Larsson L, Wu C, Fürth D, Qian X, Wärdell E, Custodio J, Reimegård J, Salmén F, et al. A Spatiotemporal Organ-Wide Gene Expression and Cell Atlas of the Developing Human Heart. Cell. 2019:179(7):1647–1660.e19. 10.1016/j.cell.2019.11.025

Backhaus AE, Griffiths C, Vergara-Cruces A, Simmonds J, Lee R, Morris RJ, and Uauy C. Delayed development of basal spikelets in wheat explains their increased floret abortion and rudimentary nature. Journal of Experimental Botany. 2023:74(17):5088–5103. 10.1093/jxb/erad233

Backhaus AE, Lister A, Tomkins M, Adamski NM, Simmonds J, Macaulay I, Morris RJ, Haerty W, and Uauy C. High expression of the MADS-box gene VRT2 increases the number of rudimentary basal spikelets in wheat. Plant Physiology. 2022:189(3):1536–1552. 10.1093/plphys/kiac156

Benfey PN and Scheres B. Root development. Current Biology. 2000:10(22):R813–R815. 10.1016/S0960-9822(00)00814-9

Benková E, Michniewicz M, Sauer M, Teichmann T, Seifertová D, Jürgens G, and Friml J. Local, Efflux-Dependent Auxin Gradients as a Common Module for Plant Organ Formation. Cell. 2003:115(5):591–602. 10.1016/S0092-8674(03)00924-3

Biancalani T, Scalia G, Buffoni L, Avasthi R, Lu Z, Sanger A, Tokcan N, Vanderburg CR, Segerstolpe Å, Zhang M, et al. Deep learning and alignment of spatially resolved single-cell transcriptomes with Tangram. Nat Methods. 2021:18(11):1352–1362. 10.1038/s41592-021-01264-7

Bradski G and Kaehler A. Learning OpenCV: Computer Vision with the OpenCV Library (O’Reilly Media, Inc.).

Cao J, Spielmann M, Qiu X, Huang X, Ibrahim DM, Hill AJ, Zhang F, Mundlos S, Christiansen L, Steemers FJ, et al. The single-cell transcriptional landscape of mammalian organogenesis. Nature. 2019:566(7745):496–502. 10.1038/s41586-019-0969-x

Chen A, Liao S, Cheng M, Ma K, Wu L, Lai Y, Qiu X, Yang J, Xu J, Hao S, et al. Spatiotemporal transcriptomic atlas of mouse organogenesis using DNA nanoball-patterned arrays. Cell. 2022:185(10):1777–1792.e21. 10.1016/j.cell.2022.04.003

Chen H, Patterson N, and Reich D. Population differentiation as a test for selective sweeps. Genome Res. 2010:20(3):393–402. 10.1101/gr.100545.109

Chen T, Chen X, Zhang S, Zhu J, Tang B, Wang A, Dong L, Zhang Z, Yu C, Sun Y, et al. The Genome Sequence Archive Family: Toward Explosive Data Growth and Diverse Data Types. Genomics Proteomics Bioinformatics. 2021:19(4):578–583. 10.1016/j.gpb.2021.08.001

Cheng S, Feng C, Wingen LU, Cheng H, Riche AB, Jiang M, Leverington-Waite M, Huang Z, Collier S, Orford S, et al. Harnessing landrace diversity empowers wheat breeding. Nature. 2024:1–3. 10.1038/s41586-024-07682-9

Chuck G, Muszynski M, Kellogg E, Hake S, and Schmidt RJ. The Control of Spikelet Meristem Identity by the branched silkless1 Gene in Maize. Science. 2002:298(5596):1238–1241. 10.1126/science.1076920

CNCB-NGDC Members and Partners. Database Resources of the National Genomics Data Center, China National Center for Bioinformation in 2022. Nucleic Acids Res. 2022:50(D1):D27–D38. 10.1093/nar/gkab951

Cucinotta M, Cavalleri A, Chandler JW, and Colombo L. Auxin and Flower Development: A Blossoming Field. Cold Spring Harb Perspect Biol. 2021:13(2):a039974. 10.1101/cshperspect.a039974

Danecek P, Auton A, Abecasis G, Albers CA, Banks E, DePristo MA, Handsaker RE, Lunter G, Marth GT, Sherry ST, et al. The variant call format and VCFtools. Bioinformatics. 2011:27(15):2156–2158. 10.1093/bioinformatics/btr330

Danecek P, Bonfield JK, Liddle J, Marshall J, Ohan V, Pollard MO, Whitwham A, Keane T, McCarthy SA, Davies RM, et al. Twelve years of SAMtools and BCFtools. Gigascience. 2021:10(2):giab008. 10.1093/gigascience/giab008

Davidson EH, Rast JP, Oliveri P, Ransick A, Calestani C, Yuh C-H, Minokawa T, Amore G, Hinman V, Arenas-Mena C, et al. A Genomic Regulatory Network for Development. Science. 2002:295(5560):1669–1678. 10.1126/science.1069883

Dobin A, Davis CA, Schlesinger F, Drenkow J, Zaleski C, Jha S, Batut P, Chaisson M, and Gingeras TR. STAR: ultrafast universal RNA-seq aligner. Bioinformatics. 2013:29(1):15–21. 10.1093/bioinformatics/bts635

Dobrovolskaya O, Pont C, Sibout R, Martinek P, Badaeva E, Murat F, Chosson A, Watanabe N, Prat E, Gautier N, et al. FRIZZY PANICLE Drives Supernumerary Spikelets in Bread Wheat1. Plant Physiol. 2015:167(1):189–199. 10.1104/pp.114.250043

Douglas SJ, Chuck G, Dengler RE, Pelecanda L, and Riggs CD. KNAT1 and ERECTA regulate inflorescence architecture in Arabidopsis. Plant Cell. 2002:14(3):547–558. 10.1105/tpc.010391

Du D, Zhang D, Yuan J, Feng M, Li Z, Wang Z, Zhang Z, Li X, Ke W, Li R, et al. FRIZZY PANICLE defines a regulatory hub for simultaneously controlling spikelet formation and awn elongation in bread wheat. New Phytologist. 2021:231(2):814–833. 10.1111/nph.17388

Fawkner-Corbett D, Antanaviciute A, Parikh K, Jagielowicz M, Gerós AS, Gupta T, Ashley N, Khamis D, Fowler D, Morrissey E, et al. Spatiotemporal analysis of human intestinal development at single-cell resolution. Cell. 2021:184(3):810–826.e23. 10.1016/j.cell.2020.12.016

Francis D. The plant cell cycle − 15 years on. New Phytologist. 2007:174(2):261–278. 10.1111/j.1469-8137.2007.02038.x

Gallavotti A, Yang Y, Schmidt RJ, and Jackson D. The Relationship between Auxin Transport and Maize Branching. Plant Physiology. 2008:147(4):1913–1923. 10.1104/pp.108.121541

Gao X-Q, Wang N, Wang X-L, and Zhang XS. Architecture of Wheat Inflorescence: Insights from Rice. Trends in Plant Science. 2019:24(9):802–809. 10.1016/j.tplants.2019.06.002

Gauley A and Boden SA. Genetic pathways controlling inflorescence architecture and development in wheat and barley. Journal of Integrative Plant Biology. 2019:61(3):296–309. 10.1111/jipb.12732

Granja JM, Corces MR, Pierce SE, Bagdatli ST, Choudhry H, Chang HY, and Greenleaf WJ. ArchR is a scalable software package for integrative single-cell chromatin accessibility analysis. Nat Genet. 2021:53(3):403–411. 10.1038/s41588-021-00790-6

Grant CE, Bailey TL, and Noble WS. FIMO: scanning for occurrences of a given motif. Bioinformatics. 2011:27(7):1017–1018. 10.1093/bioinformatics/btr064

Hama E, Takumi S, Ogihara Y, and Murai K. Pistillody is caused by alterations to the class-B MADS-box gene expression pattern in alloplasmic wheats. Planta. 2004:218(5):712–720. 10.1007/s00425-003-1157-6

Hao Y, Hao S, Andersen-Nissen E, Mauck WM, Zheng S, Butler A, Lee MJ, Wilk AJ, Darby C, Zager M, et al. Integrated analysis of multimodal single-cell data. Cell. 2021:184(13):3573–3587.e29. 10.1016/j.cell.2021.04.048

Hendelman A, Zebell S, Rodriguez-Leal D, Dukler N, Robitaille G, Wu X, Kostyun J, Tal L, Wang P, Bartlett ME, et al. Conserved pleiotropy of an ancient plant homeobox gene uncovered by *cis-*regulatory dissection. Cell. 2021:184(7):1724–1739.e16. 10.1016/j.cell.2021.02.001

Huang X, Qian Q, Liu Z, Sun H, He S, Luo D, Xia G, Chu C, Li J, and Fu X. Natural variation at the DEP1 locus enhances grain yield in rice. Nat Genet. 2009:41(4):494–497. 10.1038/ng.352

International Wheat Genome Sequencing Consortium (IWGSC). Shifting the limits in wheat research and breeding using a fully annotated reference genome. Science. 2018:361(6403):eaar7191. 10.1126/science.aar7191

Jiang D, Borg M, Lorković ZJ, Montgomery SA, Osakabe A, Yelagandula R, Axelsson E, and Berger F. The evolution and functional divergence of the histone H2B family in plants. PLOS Genetics. 2020:16(7):e1008964. 10.1371/journal.pgen.1008964

Jin J, Tian F, Yang D-C, Meng Y-Q, Kong L, Luo J, and Gao G. PlantTFDB 4.0: toward a central hub for transcription factors and regulatory interactions in plants. Nucleic Acids Research. 2017:45(D1):D1040–D1045. 10.1093/nar/gkw982

Kellogg EA. Genetic control of branching patterns in grass inflorescences. The Plant Cell. 2022:34(7):2518–2533. 10.1093/plcell/koac080

Komatsu M, Chujo A, Nagato Y, Shimamoto K, and Kyozuka J. FRIZZY PANICLE is required to prevent the formation of axillary meristems and to establish floral meristem identity in rice spikelets. Development. 2003:130(16):3841–3850. 10.1242/dev.00564

Kong X, Wang F, Geng S, Guan J, Tao S, Jia M, Sun G, Wang Z, Wang K, Ye X, et al. The wheat AGL6-like MADS-box gene is a master regulator for floral organ identity and a target for spikelet meristem development manipulation. Plant Biotechnology Journal. 2022:20(1):75–88. 10.1111/pbi.13696

Koppolu R, Chen S, and Schnurbusch T. Evolution of inflorescence branch modifications in cereal crops. Current Opinion in Plant Biology. 2022:65:102168. 10.1016/j.pbi.2021.102168

Koppolu R and Schnurbusch T. Developmental pathways for shaping spike inflorescence architecture in barley and wheat. Journal of Integrative Plant Biology. 2019:61(3):278–295. 10.1111/jipb.12771

Lareau CA, Duarte FM, Chew JG, Kartha VK, Burkett ZD, Kohlway AS, Pokholok D, Aryee MJ, Steemers FJ, Lebofsky R, et al. Droplet-based combinatorial indexing for massive-scale single-cell chromatin accessibility. Nat Biotechnol. 2019:37(8):916–924. 10.1038/s41587-019-0147-6

Lei Y, Cheng M, Li Z, Zhuang Z, Wu L, Sun Y, Han L, Huang Z, Wang Y, Wang Z, et al. Spatially resolved gene regulatory and disease-related vulnerability map of the adult Macaque cortex. Nat Commun. 2022:13(1):6747. 10.1038/s41467-022-34413-3

Li H. Aligning sequence reads, clone sequences and assembly contigs with BWA-MEM. 2013. 10.48550/arXiv.1303.3997

Li K, Debernardi JM, Li C, Lin H, Zhang C, Jernstedt J, Korff M von, Zhong J, and Dubcovsky J. Interactions between SQUAMOSA and SHORT VEGETATIVE PHASE MADS-box proteins regulate meristem transitions during wheat spike development. The Plant Cell. 2021a:33(12):3621–3644. 10.1093/plcell/koab243

Li Y, Fu X, Zhao M, Zhang W, Li B, An D, Li J, Zhang A, Liu R, and Liu X. A Genome-wide View of Transcriptome Dynamics During Early Spike Development in Bread Wheat. Sci Rep. 2018:8(1):15338. 10.1038/s41598-018-33718-y

Li Y, Li L, Zhao M, Guo L, Guo X, Zhao D, Batool A, Dong B, Xu H, Cui S, et al. Wheat FRIZZY PANICLE activates VERNALIZATION1-A and HOMEOBOX4-A to regulate spike development in wheat. Plant Biotechnology Journal. 2021b:19(6):1141– 1154. 10.1111/pbi.13535

Lin X, Xu Y, Wang D, Yang Y, Zhang X, Bie X, Gui L, Chen Z, Ding Y, Mao L, et al. Systematic identification of wheat spike developmental regulators by integrated multi-omics, transcriptional network, GWAS, and genetic analyses. Molecular Plant. 2024:17(3):438–459. 10.1016/j.molp.2024.01.010

Liu C, Wu T, Fan F, Liu Y, Wu L, Junkin M, Wang Z, Yu Y, Wang W, Wei W, et al. A portable and cost-effective microfluidic system for massively parallel single-cell transcriptome profiling. 2019:818450. 10.1101/818450

Liu J, Chen Z, Wang Z, Zhang Z, Xie X, Wang Z, Chai L, Song L, Cheng X, Feng M, et al. Ectopic expression of *VRT-A2* underlies the origin of *Triticum polonicum* and *Triticum petropavlovskyi* with long outer glumes and grains. Molecular Plant. 2021:14(9):1472–1488. 10.1016/j.molp.2021.05.021

Liu X, Bie XM, Lin X, Li M, Wang H, Zhang X, Yang Y, Zhang C, Zhang XS, and Xiao J. Uncovering the transcriptional regulatory network involved in boosting wheat regeneration and transformation. Nat Plants. 2023:9(6):908–925. 10.1038/s41477-023-01406-z

Long KA, Lister A, Jones MRW, Adamski NM, Ellis RE, Chedid C, Carpenter SJ, Liu X, Backhaus AE, Goldson A, et al. Spatial Transcriptomics Reveals Expression Gradients in Developing Wheat Inflorescences at Cellular Resolution. 2024:2024.12.19.629411. 10.1101/2024.12.19.629411

Lun ATL, Riesenfeld S, Andrews T, Dao TP, Gomes T, Marioni JC, and participants in the 1st Human Cell Atlas Jamboree. EmptyDrops: distinguishing cells from empty droplets in droplet-based single-cell RNA sequencing data. Genome Biology. 2019:20(1):63. 10.1186/s13059-019-1662-y

Luo X, Yang Y, Lin X, and Xiao J. Deciphering spike architecture formation towards yield improvement in wheat. Journal of Genetics and Genomics. 2023:50(11):835–845. 10.1016/j.jgg.2023.02.015

Marand AP, Chen Z, Gallavotti A, and Schmitz RJ. A *cis*-regulatory atlas in maize at single-cell resolution. Cell. 2021:184(11):3041–3055.e21. 10.1016/j.cell.2021.04.014

McSteen P, Laudencia-Chingcuanco D, and Colasanti J. A floret by any other name: control of meristem identity in maize. Trends in Plant Science. 2000:5(2):61–66. 10.1016/S1360-1385(99)01541-1

Moffitt JR, Bambah-Mukku D, Eichhorn SW, Vaughn E, Shekhar K, Perez JD, Rubinstein ND, Hao J, Regev A, Dulac C, et al. Molecular, spatial, and functional single-cell profiling of the hypothalamic preoptic region. Science. 2018:362(6416):eaau5324. 10.1126/science.aau5324

Neubert P and Protzel P. Compact Watershed and Preemptive SLIC: On Improving Trade-offs of Superpixel Segmentation Algorithms.. In. 2014 22nd International Conference on Pattern Recognition., pp. 996–1001. 10.1109/ICPR.2014.181

Niu J, Ma S, Zheng S, Zhang C, Lu Y, Si Y, Tian S, Shi X, Liu X, Naeem MK, et al. Whole-genome sequencing of diverse wheat accessions uncovers genetic changes during modern breeding in China and the United States. The Plant Cell. 2023:35(12):4199–4216. 10.1093/plcell/koad229

Pachitariu M and Stringer C. Cellpose 2.0: how to train your own model. Nat Methods. 2022:19(12):1634–1641. 10.1038/s41592-022-01663-4

Paolacci AR, Tanzarella OA, Porceddu E, Varotto S, and Ciaffi M. Molecular and phylogenetic analysis of MADS-box genes of MIKC type and chromosome location of SEP-like genes in wheat (Triticum aestivum L.). Mol Genet Genomics. 2007:278(6):689–708. 10.1007/s00438-007-0285-2

Pedersen DS and Grasser KD. The role of chromosomal HMGB proteins in plants. Biochimica et Biophysica Acta (BBA) - Gene Regulatory Mechanisms. 2010:1799(1):171–174. 10.1016/j.bbagrm.2009.11.004

Postma-Haarsma AD, Rueb S, Scarpella E, den Besten W, Hoge JHC, and Meijer AH. Developmental regulation and downstream effects of the knox class homeobox genes Oskn2 and Oskn3 from rice. Plant Mol Biol. 2002:48(4):423–441. 10.1023/A:1014047917226

Poursarebani N, Seidensticker T, Koppolu R, Trautewig C, Gawroński P, Bini F, Govind G, Rutten T, Sakuma S, Tagiri A, et al. The Genetic Basis of Composite Spike Form in Barley and “Miracle-Wheat.” Genetics. 2015:201(1):155–165. 10.1534/genetics.115.176628

Prusinkiewicz P, Erasmus Y, Lane B, Harder LD, and Coen E. Evolution and Development of Inflorescence Architectures. Science. 2007:316(5830):1452– 1456. 10.1126/science.1140429

Qiu X, Mao Q, Tang Y, Wang L, Chawla R, Pliner HA, and Trapnell C. Reversed graph embedding resolves complex single-cell trajectories. Nat Methods. 2017:14(10):979–982. 10.1038/nmeth.4402

Ramírez-González RH, Borrill P, Lang D, Harrington SA, Brinton J, Venturini L, Davey M, Jacobs J, van Ex F, Pasha A, et al. The transcriptional landscape of polyploid wheat. Science. 2018:361(6403):eaar6089. 10.1126/science.aar6089

Reinhardt D, Mandel T, and Kuhlemeier C. Auxin Regulates the Initiation and Radial Position of Plant Lateral Organs. The Plant Cell. 2000:12(4):507–518. 10.1105/tpc.12.4.507

Saeed S, Usman B, Shim S-H, Khan SU, Nizamuddin S, Saeed S, Shoaib Y, Jeon J-S, and Jung K-H. CRISPR/Cas-mediated editing of *cis*-regulatory elements for crop improvement. Plant Science. 2022:324:111435. 10.1016/j.plantsci.2022.111435

Sakuma S and Schnurbusch T. Of floral fortune: tinkering with the grain yield potential of cereal crops. New Phytologist. 2020:225(5):1873–1882. 10.1111/nph.16189

Sawchuk MG, Donner TJ, Head P, and Scarpella E. Unique and Overlapping Expression Patterns among Members of Photosynthesis-Associated Nuclear Gene Families in Arabidopsis. Plant Physiology. 2008:148(4):1908–1924. 10.1104/pp.108.126946

Scarpella E, Marcos D, Friml J, and Berleth T. Control of leaf vascular patterning by polar auxin transport. Genes Dev. 2006:20(8):1015–1027. 10.1101/gad.1402406

Shi Q, Liu S, Kristiansen K, and Liu L. The FASTQ+ format and PISA. Bioinformatics. 2022:38(19):4639–4642. 10.1093/bioinformatics/btac562

Shitsukawa N, Kinjo H, Takumi S, and Murai K. Heterochronic development of the floret meristem determines grain number per spikelet in diploid, tetraploid and hexaploid wheats. Ann Bot. 2009:104(2):243–251. 10.1093/aob/mcp129

Signor SA and Nuzhdin SV. The Evolution of Gene Expression in cis and trans. Trends in Genetics. 2018:34(7):532–544. 10.1016/j.tig.2018.03.007

Song X, Meng X, Guo H, Cheng Q, Jing Y, Chen M, Liu G, Wang B, Wang Y, Li J, et al. Targeting a gene regulatory element enhances rice grain yield by decoupling panicle number and size. Nat Biotechnol. 2022:40(9):1403–1411. 10.1038/s41587-022-01281-7

Sun J, Bie XM, Chu XL, Wang N, Zhang XS, and Gao X-Q. Genome-edited TaTFL1-5 mutation decreases tiller and spikelet numbers in common wheat. Frontiers in Plant Science. 2023:14.

Swinnen G, Goossens A, and Pauwels L. Lessons from Domestication: Targeting *Cis*-Regulatory Elements for Crop Improvement. Trends in Plant Science. 2016:21(6):506–515. 10.1016/j.tplants.2016.01.014

Taguchi-Shiobara F, Kawagoe Y, Kato H, Onodera H, Tagiri A, Hara N, Miyao A, Hirochika H, Kitano H, Yano M, et al. A loss-of-function mutation of rice DENSE PANICLE 1 causes semi-dwarfness and slightly increased number of spikelets. Breeding Science. 2011:61(1):17–25. 10.1270/jsbbs.61.17

Taylor-Teeples M, Lanctot A, and Nemhauser JL. As above, so below: Auxin’s role in lateral organ development. Developmental Biology. 2016:419(1):156–164. 10.1016/j.ydbio.2016.03.020

Vandepoele K, Raes J, De Veylder L, Rouzé P, Rombauts S, and Inzé D. Genome-wide analysis of core cell cycle genes in Arabidopsis. Plant Cell. 2002:14(4):903–916. 10.1105/tpc.010445

Waddington SR, Cartwright PM, and Wall PC. A Quantitative Scale of Spike Initial and Pistil Development in Barley and Wheat. Annals of Botany. 1983:51(1):119–130. 10.1093/oxfordjournals.aob.a086434

Wang C, Yang X, and Li G. Molecular Insights into Inflorescence Meristem Specification for Yield Potential in Cereal Crops. International Journal of Molecular Sciences. 2021:22(7):3508. 10.3390/ijms22073508

Wang D, Li Y, Wang H, Xu Y, Yang Y, Zhou Y, Chen Z, Zhou Y, Gui L, Guo Y, et al. Boosting wheat functional genomics via an indexed EMS mutant library of KN9204. Plant Comm. 2023:4(4). 10.1016/j.xplc.2023.100593

Wang Y, Du F, Wang J, Wang K, Tian C, Qi X, Lu F, Liu X, Ye X, and Jiao Y. Improving bread wheat yield through modulating an unselected AP2/ERF gene. Nat Plants. 2022:8(8):930–939. 10.1038/s41477-022-01197-9

Wang Y, Luo Y, Guo X, Li Y, Yan J, Shao W, Wei W, Wei X, Yang T, Chen J, et al. A spatial transcriptome map of the developing maize ear. Nat Plants. 2024:10(5):815–827. 10.1038/s41477-024-01683-2

Wang Y, Yu H, Tian C, Sajjad M, Gao C, Tong Y, Wang X, and Jiao Y. Transcriptome Association Identifies Regulators of Wheat Spike Architecture. Plant Physiol. 2017:175(2):746–757. 10.1104/pp.17.00694

Whipple CJ. Grass inflorescence architecture and evolution: the origin of novel signaling centers. New Phytologist. 2017:216(2):367–372. 10.1111/nph.14538

Xia K, Sun H-X, Li J, Li J, Zhao Y, Chen L, Qin C, Chen R, Chen Z, Liu G, et al. The single-cell stereo-seq reveals region-specific cell subtypes and transcriptome profiling in Arabidopsis leaves. Developmental Cell. 2022:57(10):1299–1310.e4. 10.1016/j.devcel.2022.04.011

Xiao J and Wagner D. Polycomb repression in the regulation of growth and development in Arabidopsis. Current Opinion in Plant Biology. 2015:23:15–24. 10.1016/j.pbi.2014.10.003

Xu X, Crow M, Rice BR, Li F, Harris B, Liu L, Demesa-Arevalo E, Lu Z, Wang L, Fox N, et al. Single-cell RNA sequencing of developing maize ears facilitates functional analysis and trait candidate gene discovery. Developmental Cell. 2021:56(4):557–568.e6. 10.1016/j.devcel.2020.12.015

Yang J, Yuan Z, Meng Q, Huang G, Périn C, Bureau C, Meunier A-C, Ingouff M, Bennett MJ, Liang W, et al. Dynamic Regulation of Auxin Response during Rice Development Revealed by Newly Established Hormone Biosensor Markers. Front Plant Sci. 2017:8. 10.3389/fpls.2017.00256

Yoshida A, Sasao M, Yasuno N, Takagi K, Daimon Y, Chen R, Yamazaki R, Tokunaga H, Kitaguchi Y, Sato Y, et al. TAWAWA1, a regulator of rice inflorescence architecture, functions through the suppression of meristem phase transition. Proc Natl Acad Sci U S A. 2013:110(2):767–772. 10.1073/pnas.1216151110

Yoshikawa GV and Boden SA. Finding the right balance: The enduring role of florigens during cereal inflorescence development and their influence on fertility. Current Opinion in Plant Biology. 2024:79:102539. 10.1016/j.pbi.2024.102539

Yu G, Wang L-G, and He Q-Y. ChIPseeker: an R/Bioconductor package for ChIP peak annotation, comparison and visualization. Bioinformatics. 2015:31(14):2382–2383. 10.1093/bioinformatics/btv145

Zhang L, He C, Lai Y, Wang Y, Kang L, Liu A, Lan C, Su H, Gao Y, Li Z, et al. Asymmetric gene expression and cell-type-specific regulatory networks in the root of bread wheat revealed by single-cell multiomics analysis. Genome Biology. 2023:24(1):65. 10.1186/s13059-023-02908-x

Zhang T-Q, Chen Y, and Wang J-W. A single-cell analysis of the *Arabidopsis* vegetative shoot apex. Developmental Cell. 2021a:56(7):1056–1074.e8. 10.1016/j.devcel.2021.02.021

Zhang X, Luo Z, Marand AP, Yan H, Jang H, Bang S, Mendieta JP, Minow MAA, and Schmitz RJ. A spatially resolved multi-omic single-cell atlas of soybean development. Cell. 2024a:0(0). 10.1016/j.cell.2024.10.050

Zhang X, Meng W, Liu D, Pan D, Yang Y, Chen Z, Ma X, Yin W, Niu M, Dong N, et al. Enhancing rice panicle branching and grain yield through tissue-specific brassinosteroid inhibition. Science. 2024b:383(6687):eadk8838. 10.1126/science.adk8838

Zhang Y, Liu T, Meyer CA, Eeckhoute J, Johnson DS, Bernstein BE, Nusbaum C, Myers RM, Brown M, Li W, et al. Model-based analysis of ChIP-Seq (MACS). Genome Biol. 2008:9(9):R137. 10.1186/gb-2008-9-9-r137

Zhang Y, Mitsuda N, Yoshizumi T, Horii Y, Oshima Y, Ohme-Takagi M, Matsui M, and Kakimoto T. Two types of bHLH transcription factor determine the competence of the pericycle for lateral root initiation. Nat Plants. 2021b:7(5):633–643. 10.1038/s41477-021-00919-9

Zhu Y and Wagner D. Plant Inflorescence Architecture: The Formation, Activity, and Fate of Axillary Meristems. Cold Spring Harb Perspect Biol. 2020:12(1):a034652. 10.1101/cshperspect.a034652

Zong J, Wang L, Zhu L, Bian L, Zhang B, Chen X, Huang G, Zhang X, Fan J, Cao L, et al. A rice single cell transcriptomic atlas defines the developmental trajectories of rice floret and inflorescence meristems. New Phytologist. 2022:234(2):494–512. 10.1111/nph.18008

